# Mapping and Modeling Age-Related Changes in Intrinsic Neural Timescales

**DOI:** 10.1101/2024.09.10.612380

**Authors:** Kaichao Wu, Leonardo L. Gollo

**Affiliations:** Brain Networks and Modelling Laboratory and The Turner Institute for Brain and Mental Health, Monash University, 18 Innovation Walk, Melbourne, 3800, VIC, Australia; Monash Biomedical imaging, Monash University, 770 Blackburn Road, Melbourne, 3800, VIC, Australia; Instituto de Física Interdisciplinar y Sistemas Complejos, IFISC (UIB-CSIC), Campus Universitat de les Illes Balears 07122, Palma de Mallorca, Spain

**Keywords:** Aging, Criticality, Hierarchy of timescales, Neuronal network modeling, Mnemonic discrimination ability

## Abstract

Intrinsic timescales of brain regions exhibit heterogeneity, escalating with hierarchical levels, and are crucial for the temporal integration of external stimuli. Normal aging, often associated with cognitive deficits, involves progressive neuronal and synaptic loss, shaping brain structure and dynamics. The impact of these structural changes on temporal integration coding in the aged brain remains largely unknown. To address this gap, we mapped differences in intrinsic timescales and gray matter volume (GMV) within brain regions using magnetic resonance imaging (MRI) in young and elderly adults. We found shorter intrinsic timescales across multiple large-scale functional networks in the elderly cohort, and a significant positive association between intrinsic timescales and GMV. Additionally, age-related decline in performance on a visual discrimination task was linked to a reduction in intrinsic timescales in the cuneus. To explain these age-related changes in GMV and intrinsic timescales, we propose an age-dependent model of spiking neuron networks. In younger subjects, brain regions were modeled in a near-critical branching regime, while in elderly subjects, regions had fewer neurons and synapses, pushing the dynamics toward a more subcritical regime. The empirical results were reproduced by the model: Neuronal networks representing brain regions in young subjects exhibited longer intrinsic timescales due to critical slowing down. These findings reveal how age-related structural brain changes may drive alterations in brain dynamics, offering testable predictions and informing future interventions for cognitive decline.

## 1 Introduction

The brain can be conceptualized as a dynamic and evolving multiscale network, distinguished by its intricate topology and dynamic processes, operating across various spatial and temporal scales [1]. Across the lifespan, both the structural architecture and functional dynamics of the brain [2] demonstrate remarkable adaptability and plasticity [3]. Brain plasticity encompasses a diverse array of processes essential for supporting the network’s continuous evolution. These processes actively shape the structural and functional landscape of the brain [4, 5], regulating its response to internal and external stimuli, ranging from rapid synaptic adaptations occurring within milliseconds to the gradual effects of aging spanning years or decades [3]. Within this framework, the phenomenon of healthy aging emerges as a result of countless plasticity processes.

Aging exerts profound effects on the brain [6], inducing alterations across multiple spatial scales [7] that affect cognitive, sensory, and motor functions [8–10]. At the microscale, aging is marked by progressive neuronal loss, structural deterioration of dendrites and synapses [6, 11–14], and reduced neuronal firing rate [8, 14]. At the macroscale, both structural and functional connectivity undergo significant transformations [2, 4, 15]. Despite substantial age-related changes in brain structure and dynamics across scales [12, 16], fundamental features of brain function persist throughout the lifespan [17]. The brain’s capacity to adapt to a variety of stimuli demands a flexible network architecture, which is supported by a hierarchy of timescales [18– 21]. This hierarchy of timescales serves as an organizational principle that facilitates efficient processing and integration of information [22–24], particularly in dynamically changing environments [18, 20]. Moreover, the heterogeneity across brain regions, as dictated by the hierarchy of timescales, enhances the functionality of the brain by enabling specialized processing and integration of information [23].

Intrinsic timescales reflect regional heterogeneity [19, 24, 25] and key aspects of brain function and dysfunction [26]. Imbalances in intrinsic timescales within specific brain regions have been implicated in a wide range of neurological and neuropsychiatric disorders, including autism spectrum disorder [27], psychosis [28], depression [29, 30], epilepsy [31, 32], schizophrenia [33, 34], Parkinson’s disease [35], and Alzheimer’s disease [36]. This fast-growing body of research illustrates a significant interest in intrinsic timescales. However, intrinsic timescales are rarely investigated in the context of aging. Understanding how intrinsic timescales are influenced by aging can provide deeper insights into the mechanisms that underpin both healthy cognitive aging and the development of age-related neurological and neuropsychiatric disorders [27, 37– 39]. Furthermore, little evidence exists regarding the underlying mechanisms involved in imbalanced intrinsic timescales and their behavioral implications.

Intrinsic neural timescales are operationally measured based on the decay properties of the autocorrelation function [19, 24, 27, 38, 40]. The autocorrelation function quantifies how neural activity at one time point relates to the activity at subsequent time points, and maximum temporal autocorrelation occurs at criticality [41], meaning that systems approaching a critical state take longer to return to equilibrium [42]. This phenomenon of the slow decay of the autocorrelation function is known in the literature as critical slowing down [41–47]. Systems in a critical point are also associated with several optimal computational properties [41, 48–51], including a maximum number of metastable states and dynamical repertoire [42, 52], and maximum dynamic range [53–55].

Criticality can offer valuable insights into the dynamics of neural systems, helping to mechanistically explain features observed in intrinsic neural timescales often identified through empirical studies. Within this framework, a reduction in intrinsic timescales implies that the system is deviating from a critical point [22, 49, 56]. This shift could result in diminished computational capabilities and altered information processing, reflecting changes in the structural and functional integrity of neural networks.

Utilizing the theoretical framework of criticality, we study the impact of aging on brain structure and function by examining neuronal network dynamics, gray matter volume, and intrinsic timescales in young and elderly adults. We employ functional magnetic resonance imaging (fMRI) from the two cohorts to map intrinsic timescales and structural MRI to explore potential neuronal substrates by investigating gray matter volume (GMV), focusing on disparities between the two groups. By examining the dynamics of recognized resting-state functional networks, we uncover variations in intrinsic neural timescales linked to age and mnemonic discrimination ability. Furthermore, as a proof-of-principle, we utilize a parsimonious neuron network model, guided by our empirical findings on gray matter volume, to offer mechanistic insights into how aging impacts brain network structure, dynamics, and intrinsic timescales.

## 2 Results

We investigate intrinsic timescales in the human brain, extracted from fMRI data, and examine their underlying neural mechanisms through neuronal network modeling. Figure 1 summarizes the main steps used to process structural and functional MRI data, and to model neuronal network activity representing brain regions in young (18–32 years old, mean: 22.21, SD: 3.65) and elderly (61–80 years old, mean: 69.82, SD: 5.64) adults. The activated functional regions in the two cohorts were identified using group-independent component analysis (gICA) with resting-state fMRI. Both cohorts involve normal, healthy subjects with no cognitive or functional impairment. The intrinsic timescales of brain functional regions were then estimated as the area under the autocorrelation function (see Figure 1**A**, and subsection 4.5 for a thorough description). Next, we explored the neuroanatomic substrates of age-induced alterations in intrinsic timescales by mapping the GMV of the corresponding functional regions obtained from structural MRI (Figure 1**B**).

**Fig. 1.**
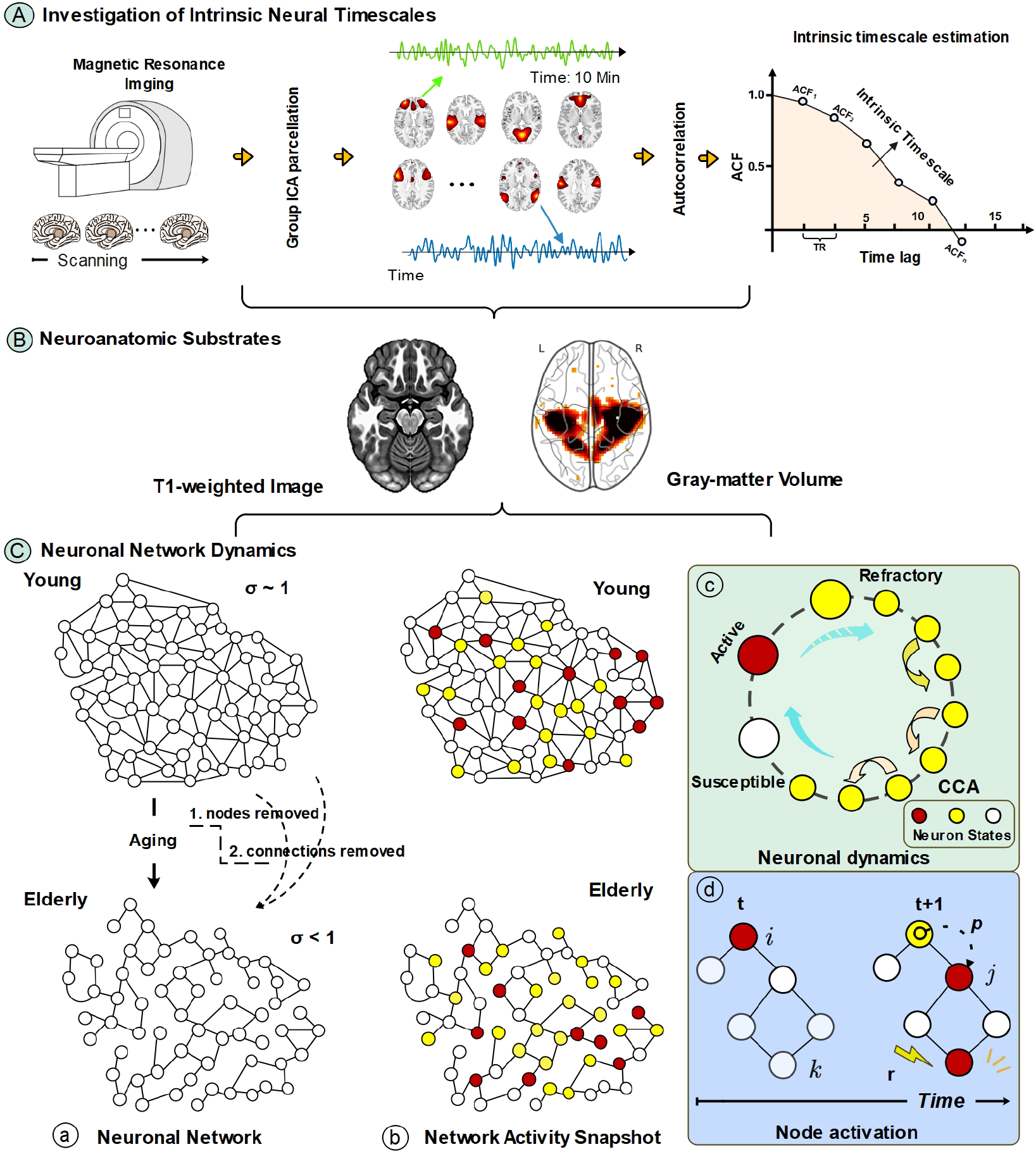
Mapping and modeling the impact of age on intrinsic neural timescales. **A**. Intrinsic timescales in the brain of younger and older adults were estimated with fMRI. Group-independent component analysis (gICA) was applied to functional images of the two cohorts to detect resting-state networks (RSN), and intrinsic timescales were calculated as the area under the autocorrelation curve of the extracted timeseries. **B**. Gray matter volume (GMV) was investigated using structural T1-weighed images to explore neuroanatomic substrates of intrinsic timescales. **C**. Neuronal network dynamics representing brain regions were examined for young and elderly adults. The aging process was depicted as neuronal and synaptic removal **(a)**. The Kinouchi-Copelli model [53] was used to simulate the dynamics of excitable neuronal networks **(b)**. The neuronal dynamics was modeled as a cyclic cellular automaton (CCA). A CCA of neurons is a discrete-time process defined on the state space where, as time evolves, neurons transit between three states: susceptible, active, and refractory **(c)**. Susceptible neurons can become active by receiving an external stimulus or a spike coming from active neighboring neurons **(d)**. Additional details can be found in the Methods section.

We employed a parsimonious neuronal network modeling approach to investigate how intrinsic timescales vary with age. (Figure 1**C**). The neuronal network dynamics is based on the Kinouchi-Copelli model [53], which clearly distinguishes between different types of branching processes: subcritical, critical, and supercritical. This influential approach considers a random network of neurons with excitable dynamics, and the branching process is controlled by the branching ratio *σ* [57]. For *σ* < 1, the system is subcritical, and each spiking neuron on average generates less than one activation, causing the network activity to decay. For *σ* = 1, the system is critical, and each spiking neuron on average generates one activation, maintaining network activity bounded and stable, despite the large fluctuations typically observed at the critical state. For *σ* > 1, the system is supercritical, and each spiking neuron on average generates more than one activation, causing the network activity to grow. In the mean-field approximation, *σ* = *K* ·*λ*, where *K* is the mean network degree, and *λ* is the probability of propagation of a neuronal spike to a neighbor susceptible neuron. Brain regions were modeled as large networks of spiking neurons. Due to the aging process, brain networks representing elderly subjects were modelled with a reduced number of neurons and synapses compared to networks representing young subjects.

### 2.1 Intrinsic Timescales for Elderly and Young Cohorts

MRI scans recorded at the University of North Carolina [58] were used comprising 34 young (20 females) and 28 elderly adults (20 females). The resting-state fMRI scans have 300 volumes (600s) of valid signals, with a repetition time (TR) of 2s, and the first five volumes (10s) were removed for magnetization stability. The functional images were pre-processed and corrected for head motion. As shown in Figure 2, resting-state network (RSN) components were computed using the spatial group independent components analysis (gICA) based on the components estimated from resting-state functional MRI data of 405 healthy controls and previous studies [59]. The spatial map of 60 recognized RSNs can be seen in Figure 2**A**, where they are assigned to 6 functional domains/networks: The default mode network (DMN), sensorimotor network (SMN), visual network (VIS), subcortical network (SC), cognitive control network (CC), and auditory network (AUD). The spatial map of the individual RSN has been sorted according to their functional network, and full details including their coordinates can be found in Supplementary Material, Figure 1 and Table I. The spatial location of the peak coordinates of these RSNs with a radius equal to 5mm can be seen in Figure 2**B**.

**Fig. 2.**
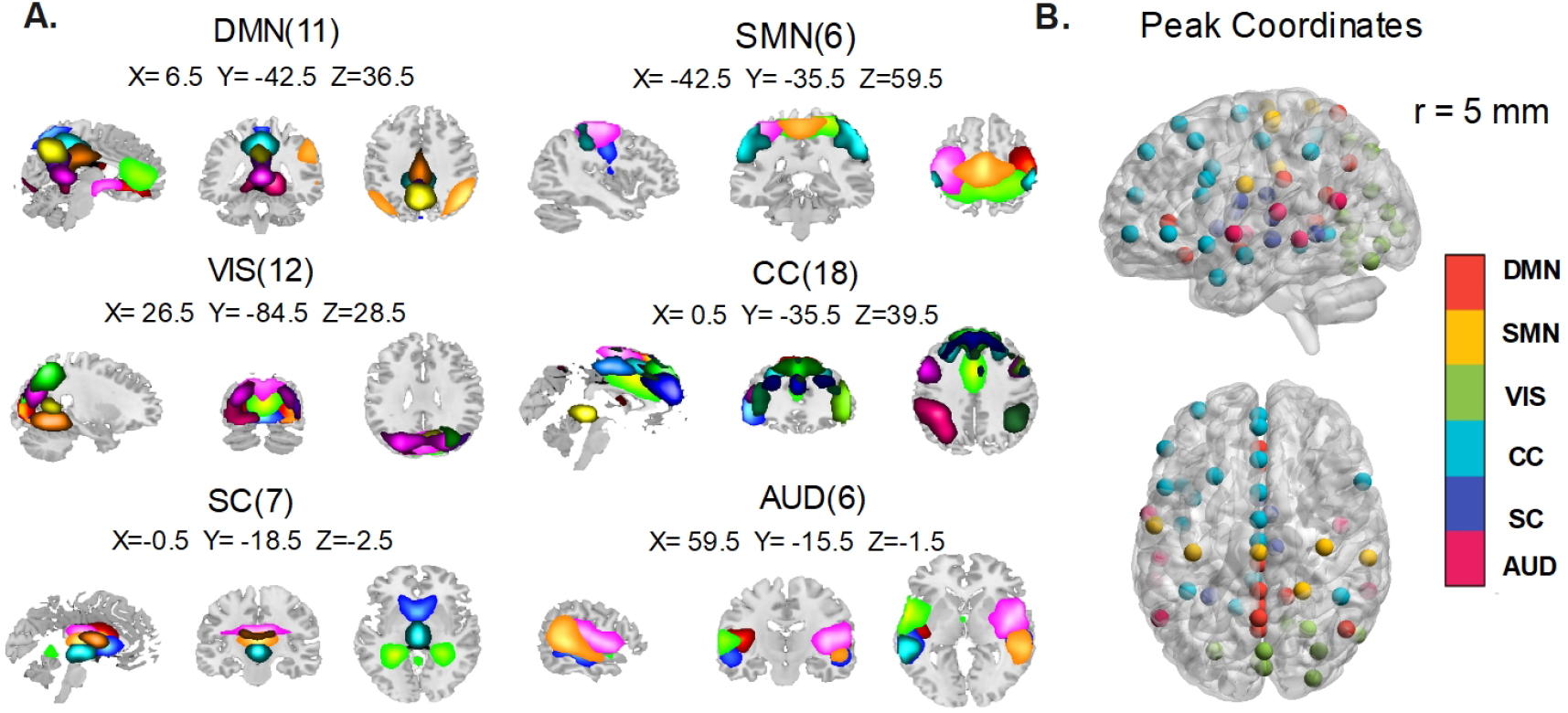
Resting-state networks. **A**. Spatial map of 60 RSN. They were assigned to six functional domains: The default mode domain (DMN), sensorimotor network (SMN), visual network (VIS), subcortical network (SC), cognitive control network (CC), and auditory network (AUD). **B**. Peak coordinates of 60 components with a radius = 5 mm and their color-coded functional networks.

For the time course of each RSN component, the intrinsic timescale was defined as the area under the curve of the autocorrelation function (see Methods). A map of intrinsic timescales of the whole brain was computed for each participant. In a comparison among functional networks, we found that both young (Figure 3**A**) and elderly (Figure 3**B**) groups show a similar hierarchy of intrinsic timescales (*F*_*young*_ = 14.612, *p* < 0.001, *F*_*elderly*_ = 15.78, *p* < 0.001, one-way ANOVA analysis), where the sub-cortical network exhibited shorter intrinsic timescales compared to other high-level cortical networks (DMN, SMN, VIS, CC, and AUD, *p* < 0.05 FDR corrected). A similar variance between the cohorts, either at a single RSN component level (Figure 3**C**) or at the network level (Figure 3**D**), shows that INT has a similar range of variability and suggests that a hierarchy of timescales plays a similar role regardless of age.

**Fig. 3.**
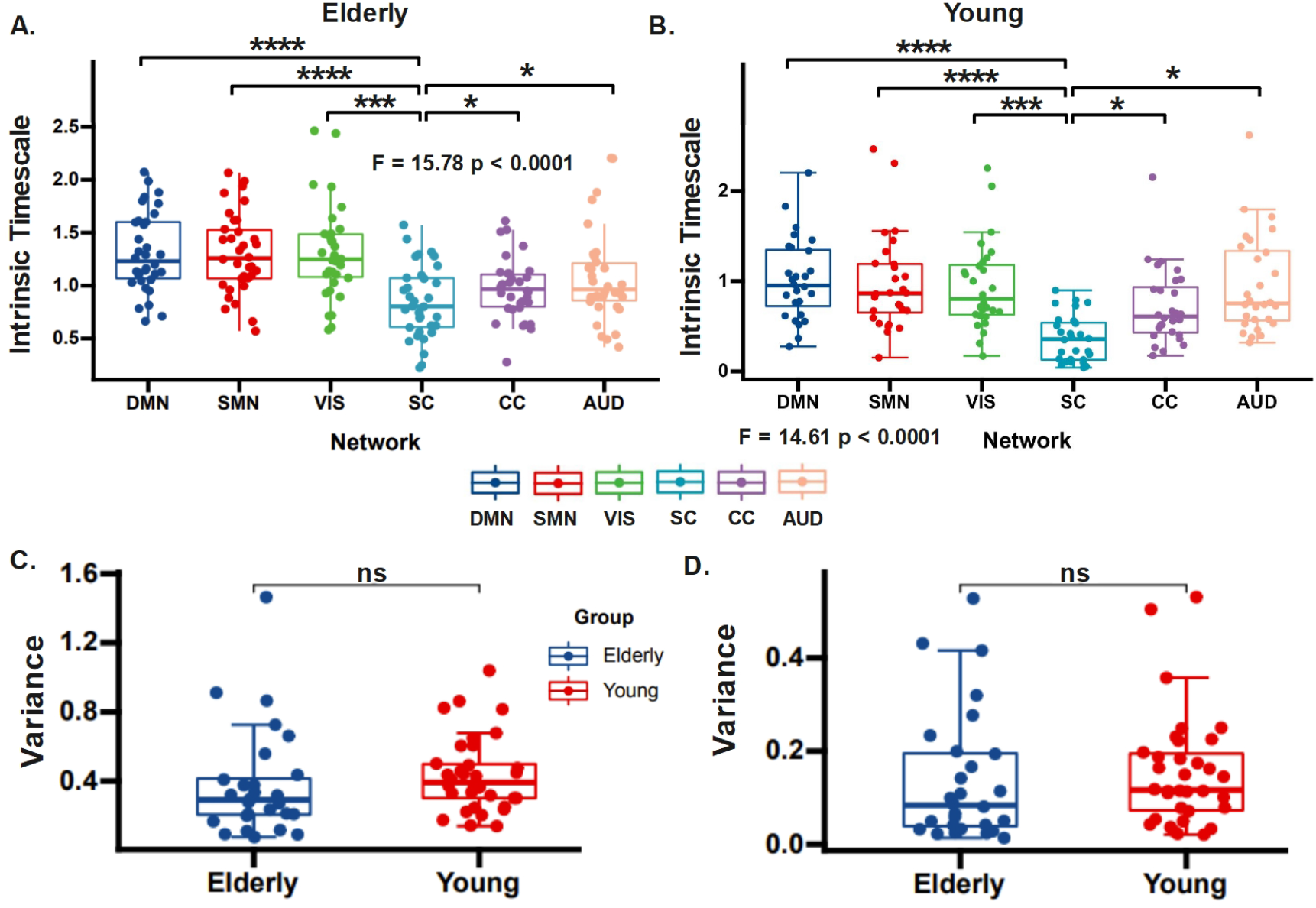
Hierarchy of timescales for young and elderly adults. A hierarchy of intrinsic timescales along the subcortical-cortical axis can be observed in both elderly **(A)** and young adults **(B)**. The variance of intrinsic timescale does not differ between the two groups both at the level of individual RSN components **(C)** and at the network level **(D)**. Boxes represent the interquartile range (IR) between the 25th and 75th percentiles. The thick line in the center of each box represents the median. The upper and lower error bars display the largest and smallest values within 1.5 times IR above the 75th percentile and below the 25th percentile, respectively. * * * indicates *p* < 0.001 (FDR-corrected); * indicates *p* < 0.05 FDR-corrected; *ns* indicates no significance.

However, the group comparison demonstrates a significant difference between the young and elderly groups in terms of intrinsic timescales of the resting-state networks. Compared with the younger cohort, older adults exhibit an overall reduction in intrinsic timescales (Welch’s t-test, *t* = 2.72, *p* < 0.001, Figure 4**A**).

**Fig. 4.**
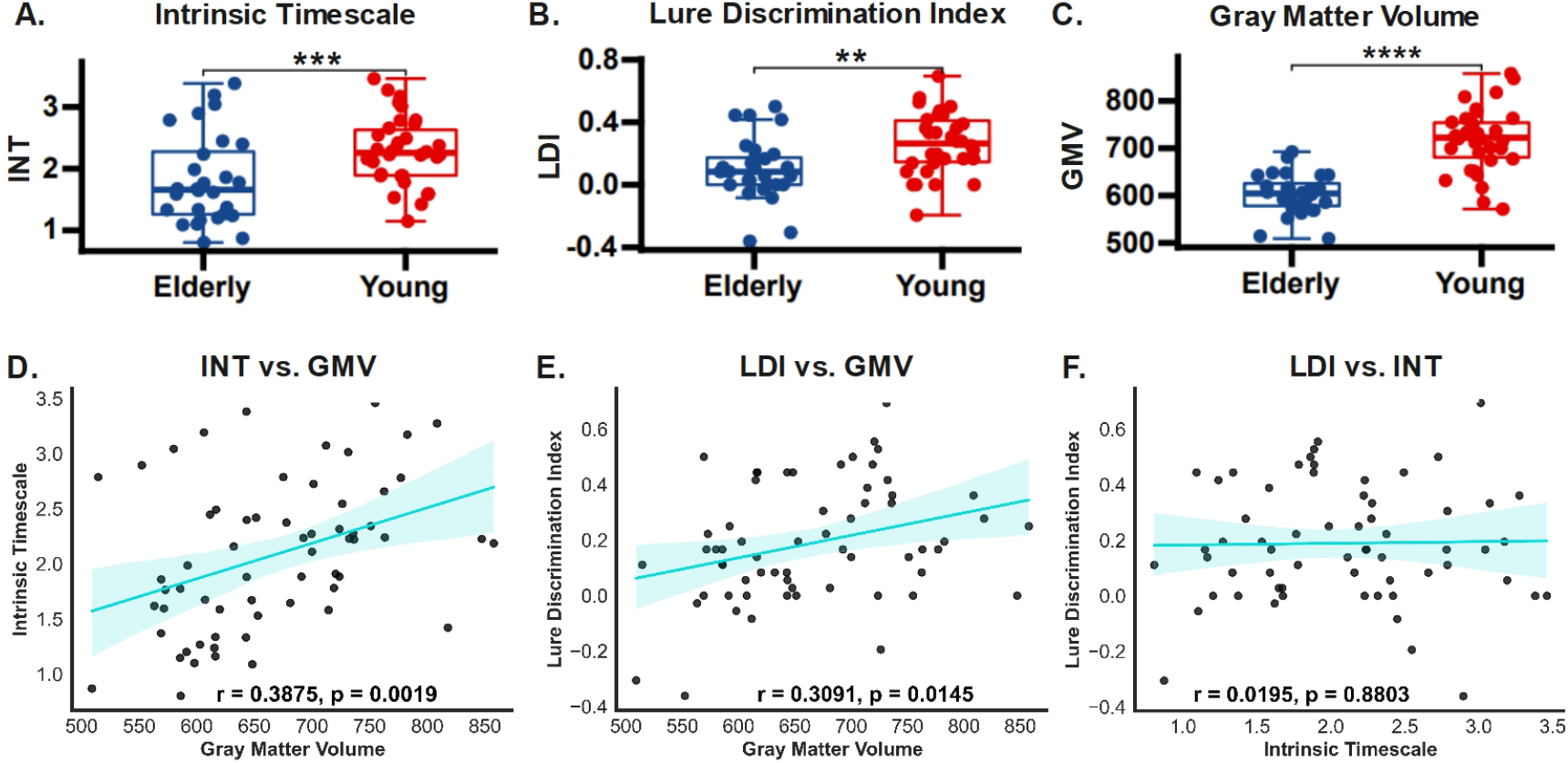
Effects of aging on brain structure, dynamics, and behavior. Comparison of intrinsic timescales (INT, **A**), lure discrimination index (LDI, **B**), and gray matter volume (GMV, **C**) between elderly and young adults. Boxes represent the interquartile range (IR) between the 25th and 75th percentiles. The thick line in the center of each box represents the median. The upper and lower error bars display the largest and smallest values within 1.5 times IR above the 75th percentile and below the 25th percentile, respectively. * * ** indicates *p* < 0.0001 (FDR-corrected); * * * indicates *p* < 0.001 (FDR-corrected); ** indicates *p* < 0.01 (FDR-corrected). Significant positive correlations were observed between GMV and INT (**D**), as well as between GMV and LDI (**E**). No association was found between INT and LDI (**F**).

### 2.2 Association Between Intrinsic Timescales, Mnemonic Discrimination Ability, and GMV

To examine the relationship between intrinsic timescales and behavior, we focus on the mnemonic discrimination ability (MDA). MDA refers to the perception capacity of an individual to accurately distinguish between similar or overlapping memories based on subtle differences in temporal context or other features. This ability can be considered essential for daily cognitive functions, and is known to decline with age [58]. MDA can be estimated by the lure discrimination index (LDI), which is calculated as the probability of the difference in response to lures and foils in a visual mnemonic discrimination task. A better MDA for the younger cohort is demonstrated by a significantly larger LDI for the young cohort compared to the elderly cohort (Welch’s t-test, *t* = 2.33, *p* < 0.05, Figure 4**B**).

To reveal the anatomic basis supporting the observed differences between groups in neural dynamics and behavior, we extracted GMV using peak coordinates of the identified RSN with a radius equal to 5mm. Similar results are obtained with alternative radius (see Supplementary Material, Figure 2). A significant difference in GMV between the two cohorts (*t*_*young−old*_ = 4.57, *p* < 0.0001) indicates that structural brain differences occur with age (Figure 4**C**). A comparison for whole-brain GMV between young and elderly subjects can be found in Supplementary Material, Figure 3. To better understand the impact of GMV, we then calculated its Person’s correlation with intrinsic timescales, and LDI. At the global level, GMV exhibited a significant positive correlation with both intrinsic timescales and LDI (INT: *r* = 0.3875, *p* < 0.05, Figure 4**D**; LDI: *r* = 0.3091, *p* < 0.05, Figure 4**E**). Nevertheless, at the whole brain average level, no significant correlation was observed between intrinsic timescales and LDI (*r* = 0.07, *p* = 0.57, Figure 4**F**). Thus, we sought to investigate intrinsic timescales at specific and specialized functional regions associated with MDA. At the single RSN level, the cuneus in the visual functional network (Figure 5**A**) shows a positive correlation between intrinsic timescales and LDI (*r* = 0.2532, *p* < 0.05, FDR corrected), and between GMV and INT (*r* = 0.3082, *p* < 0.05, FDR corrected, Figure 5**B**; see also Supplementary Material, Figure 2A).

**Fig. 5.**
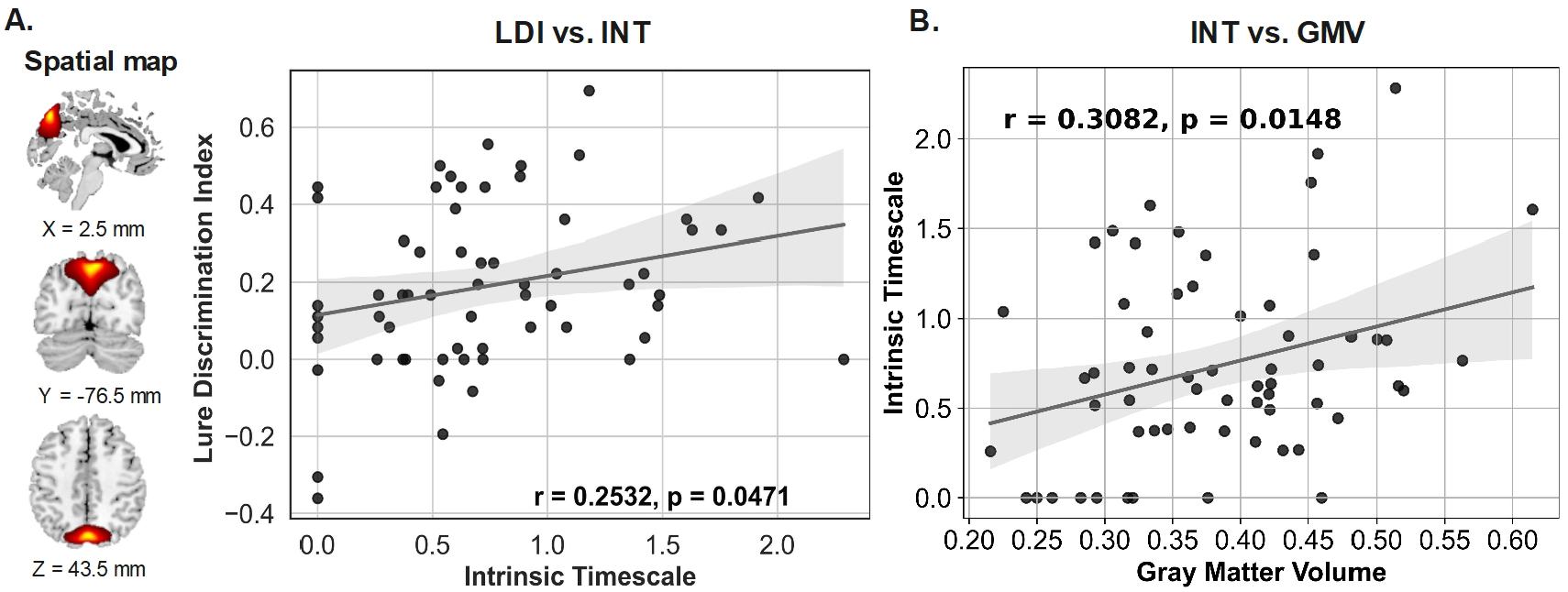
Cuneus exhibits specialized brain function. Spatial map of cuneus (A). Intrinsic timescales were found significantly correlated with LDI and GMV within the cuneus (B).

### 2.3 Relationship Between Intrinsic Timescales and GMV at the Network Level

The gICA analysis identified six large-scale function domains/networks: DMN, VIS, SMN, VIS, CC, SC, and AUD. Consistent with the whole-brain trends (Figure 4**D**), we also identified significant differences between the two groups at the network level with respect to intrinsic timescales and GMV (*p* < 0.05, FDR corrected). The elderly cohort had shorter intrinsic timescales and smaller GMV than the young cohort for all six functional networks (Figure 6**A&B**). The significantly positive correlations between GMV and INT were also robustly observed at the network level (*r*_*DMN*_ = 0.4530, *p* < 0.001; *r*_*SMN*_ = 0.4778, *p* < 0.0001; *r*_*V IS*_ = 0.3010, *p* < 0.05; *r*_*CC*_ = 0.4650, *p* < 0.001; *r*_*SC*_ = 0.3559, *p* < 0.001; *r*_*AUD*_ = 0.4445, *p* < 0.001). Results with GMV extracted with an alternative radius (= 3mm) also supports this robust association with intrinsic timescales at the network level (see Supplementary Material Figure 2B&C). These associations are consistent and reliable, remaining significant across all functional networks (Figure 6) as well as at the whole-brain level (Figure 4**D**). However, despite their reliability and significance, the interpretation of these associations is limited because they do not establish causal relationships.

**Fig. 6.**
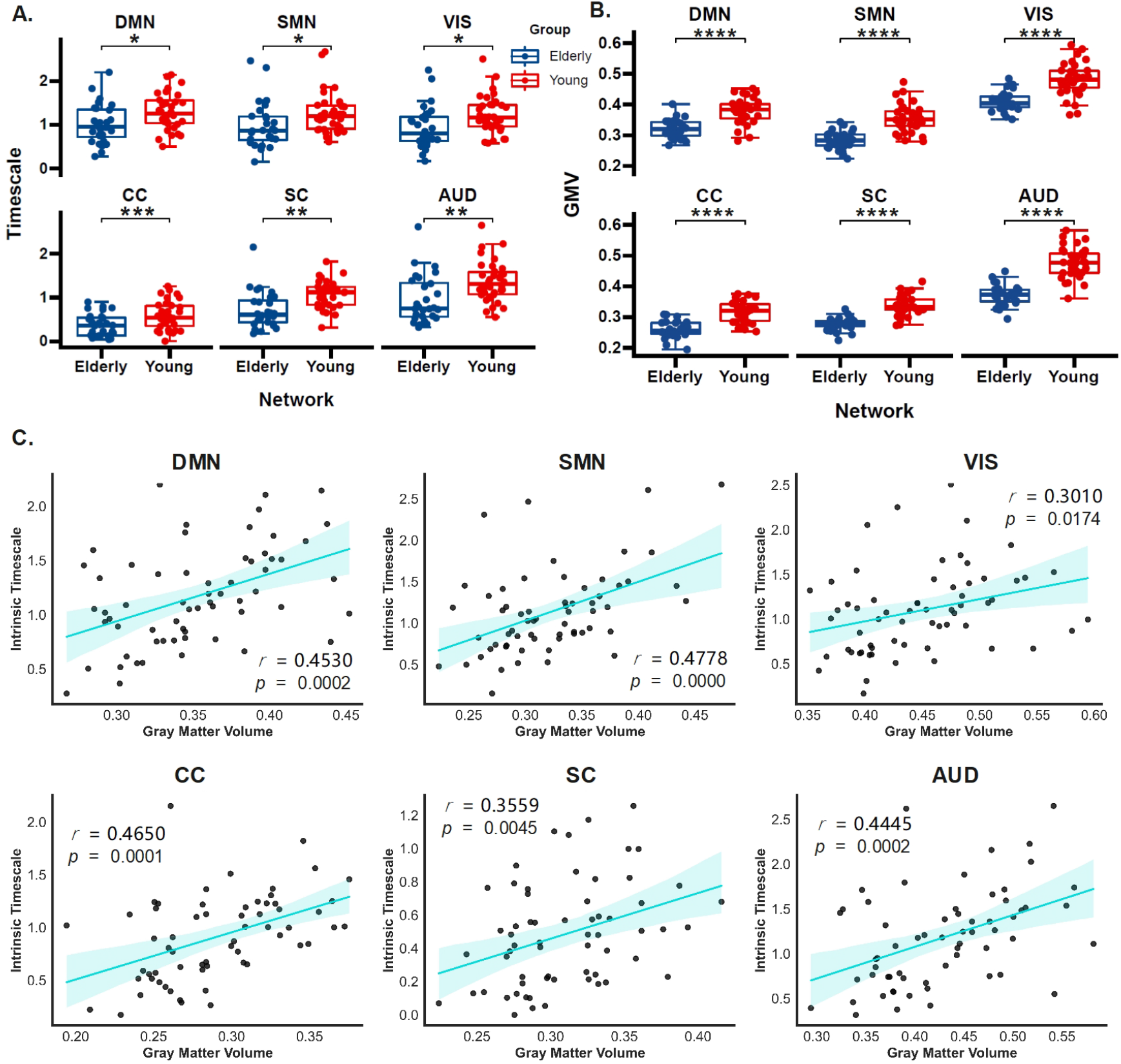
Relationship between GMV and intrinsic timescales. Comparison of intrinsic timescales **(A)** and GMV **(B)** between elderly and young adults at the network level. Positive correlations between intrinsic timescales and GMV were evident for all six functional networks**(C)**. Boxes represent the interquartile range (IR) between the 25th and 75th percentiles. The thick line in the center of each box represents the median. The upper and lower error bars display the largest and smallest values within 1.5 times IR above the 75th percentile and below the 25th percentile, respectively. * * ** indicates *p* < 0.0001 (FDR-corrected); *** indicates *p* < 0.001 (FDR-corrected); ** indicates *p* < 0.01 (FDR-corrected); * indicates *p* < 0.05 (FDR-corrected); *ns* indicates no significance.

### 2.4 Modeling Reduced Intrinsic Timescales in the Elderly

To enhance our understanding of brain dynamics and the influence of GMV on intrinsic timescales, we propose investigating a parsimonious neuronal network model that captures the dynamics of a brain region for different values of branching ratio *σ*. The strong association between GMV and intrinsic timescales suggests a potential neuroanatomical basis for the brain’s temporal dynamics. One possible interpretation for this association is that larger GMV reflects a more densely connected neuron network, where recurrent neural activity persists longer, leading to extended intrinsic timescales.

To test this hypothesis, we propose modeling brain networks for both young and elderly cohorts. Given that GMV decreases with aging, neuronal networks in the elderly should account for the loss of neurons [60, 61] and synaptic connections [62]. Reflecting the normal aging process, brain regions representing the elderly cohort are expected to exhibit a reduced number of neurons and synapses compared to those in the young cohort. To better understand the impact of aging on the dynamics of brain networks, we constructed neuronal networks representing brain regions with nearly 100,000 neurons, connected as Erdős–Rényi random networks, and simulated their dynamics. Each neuron was connected to an average of 10 other neurons via synaptic connections. Neurons and synapses were subsequently removed to simulate the aging process (see schematic representation in Figure 1 **Panel C**. a). The differences in node degree between the networks denoting the young and the elderly cohorts are illustrated in Figure 7**A**. For comparison, schematic random networks with reduced number of neurons and synapses for the elderly cohort are also displayed in Figure 7**B** to highlight the construction-imposed differences between the two networks.

**Fig. 7.**
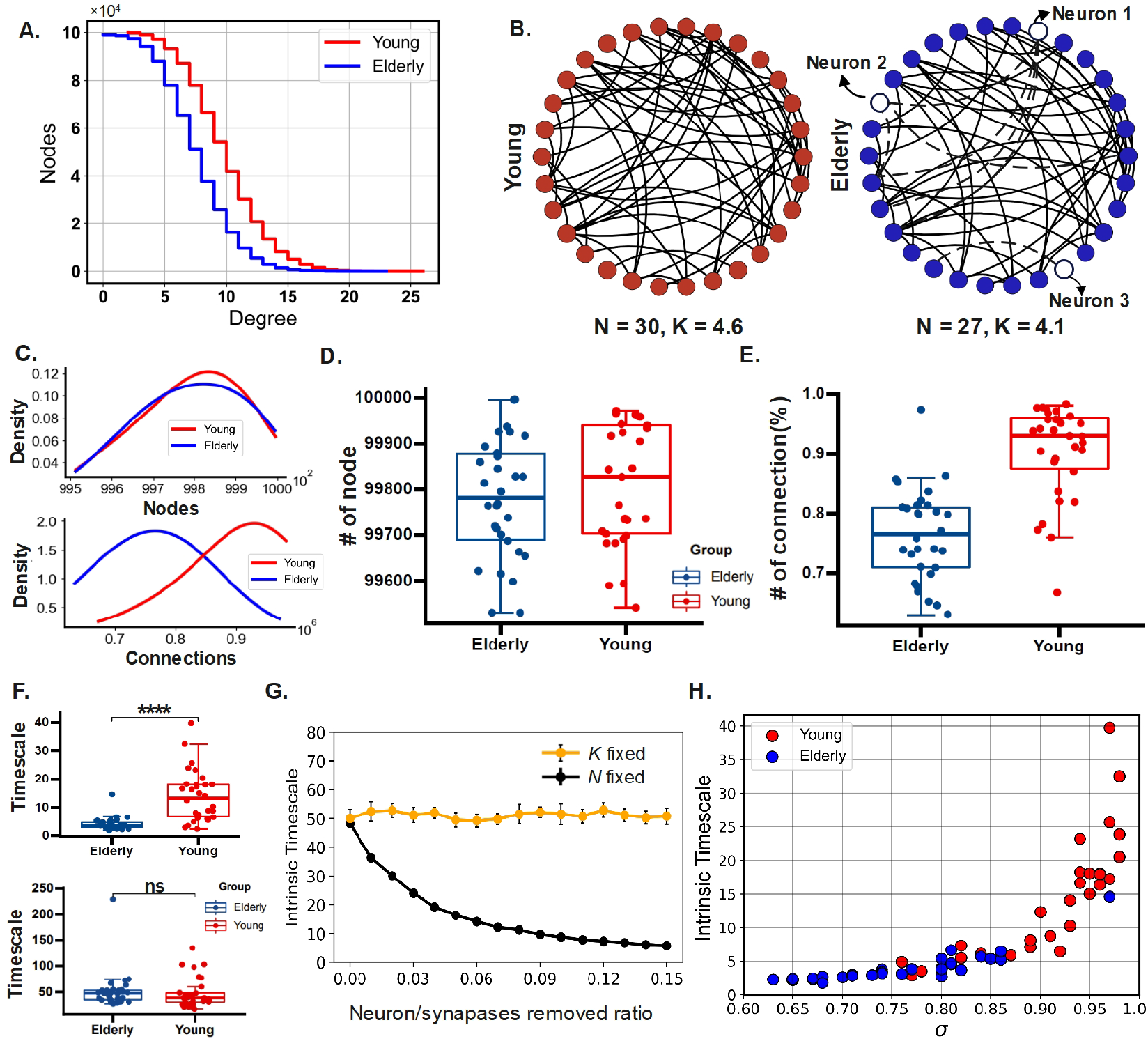
Modeling age-dependent brain networks. (A) Nodes sorted by degree in neuronal networks constructed for elderly and young adults. (B) Schematic networks illustrating age-induced differences in network structure. Left, a schematic network for a young adult (in red) has *N* = 30 neurons and an average degree *K* = 4.6. Right, a schematic network for an elderly adult (in blue) has fewer neurons and connections (*N* = 27 neurons and average degree *K* = 4.1). (C) The number of neurons (*N*) and synapses (*N*_*K*_) of 200 neuronal networks (100 for each cohort) follow a normal distribution. (D&E) Number of neurons and connections of randomly selected 60 neuronal networks, 30 for the young and 30 for the elderly cohort. (F) Intrinsic timescales calculated from neuronal network dynamics are significantly differed between the two groups only after connection removal (**top**), while removing solely neurons did not significantly alter intrinsic timescales (**bottom**). Boxes represent the interquartile range (IR) between the 25th and 75th percentiles. The thick line in the center of each box represents the median. The upper and lower error bars display the largest and smallest values within 1.5 times IR above the 75th percentile and below the 25th percentile, respectively. * * ** indicates *p* < 0.0001 (FDR-corrected); *ns* represents no significance. (G) Intrinsic timescales after systematically varying the number of neurons (with *K* = 10) or connections unilaterally (with *N* = 99, 795). (H) Intrinsic timescales with advancing age can be explained by network criticality. The younger cohort is closer to the critical point (*σ*_*c*_ = 1) than the elderly, resulting in longer intrinsic timescales.

Guided by the relative differences in GMV and INT between the two cohorts, we assumed that the number of neurons and connections removed from networks follows a normal distribution, see Methods (section 4) for details. We ran 200 trials, 100 for each cohort, to obtain distributions of the number of neurons and connections, which can be seen in Figure 7**C**. To align with the size of the empirical data, we randomly selected 30 samples representing the young cohort and 30 representing the elderly cohort for subsequent analysis. The mean number of neurons in the sample was 99812 for the young group, and 99779 for the elderly group (Figure 7**D**). The mean number of synapses was 9.15 × 10^5^ for the young group and 7.38 × 10^5^ for the elderly group (Figure 7**E**). In this sample, the ratio of the number of neurons and synapses between the young and elderly adults are *N*_*e*_*/N*_*y*_ ≈ 0.99 and *N*_*K,e*_*/N*_*K,y*_ ≈ 0.81, respectively.

Using these neuronal networks for young and elderly individuals, we simulated the networks of interacting excitable neurons with the Kinouchi-Copelli model [53]. In this model, quiescent (or susceptible) neurons spike either in response to external input *h* or through the propagation of activity from spiking neurons with probability *λ* (see Methods subsection 4.4 and Figure 1**C.c & C.d** for additional details on the neuronal dynamics). The density of spiking neurons at a given time *t* reflects the network’s instantaneous activity, which is used to compute intrinsic timescales (see Methods subsection 4.5). Consistent with the empirical data, the intrinsic timescale obtained from the network dynamics reveals a significant difference between groups (Figure 7**F** top). The elderly cohort shows substantially shorter intrinsic timescales compared to the young cohort (Welch’s t-test, *t* = 5.575, *p* < 0.0001), while removing solely neurons did not significantly alter the intrinsic timescales (Figure 7**F** bottom). Similar results were obtained after reselecting 50 and 100 samples for both young and elderly groups, showing that these findings are free from potential biases due to sample size (see Supplementary Material, Figure 4).

To further understand the effect of removing neurons and synapses on intrinsic timescales, we computed intrinsic timescales while keeping the neuronal network fixed with *K* = 10 and *N* = 99, 795 and then removing a proportion of neurons and synapses. By systematically varying the number of neurons or connections unilaterally, it is clear that removing only synapses produced a strong reduction in intrinsic timescales whilst removing only neurons did not alter them (Figure 7**G**). This finding indicates that synaptic loss has a greater impact on network dynamics than neuronal loss, confirming the significant between-group difference in intrinsic timescales (Figure 7**F**). Together, these results suggest that the reduced intrinsic timescales in elderly subjects can be caused by reduced synaptic connectivity.

### 2.5 Distance to Criticality Explains Differences in Intrinsic Timescales

Lastly, we investigated why neuronal networks with smaller GMV due to aging exhibit shorter intrinsic timescales. To address this, we focused on the distance to criticality (DTC) of the neuronal networks for both young and elderly individuals using the network branching ratio *σ*. This ratio indicates how network activity propagates [52, 53, 63]. A network in the critical state has a branching ratio *σ* = 1 and a subcritical network has *σ* < 1 [41, 50, 53, 64, 65]. For subcritical networks, a smaller value of *σ* indicates a greater DTC.

Driven by differences in network degree (Figure 7**E**), the branching ratio of neuronal networks in the younger brain was significantly higher than in the older brain (*p* < 0.001). Additionally, the relationship between intrinsic timescales and the branching ratio (Figure 7**H**) revealed that intrinsic timescales increase sharply as the network approaches the critical point (*σ* = 1). This behavior strongly indicates critical slowing down [41, 43–47]. As a result, the neuronal network in the young brain with higher intrinsic timescales (red nodes, Figure 7**H**) have a shorter DTC than those in the elderly brain (blue nodes, Figure 7**H**).

To quantitatively assess brain criticality and measure DTC for young and elderly adults, we used the maximized autocorrelation timescales, i.e., the lag-1 autocorrelation (AC1), which is regarded as a universal signature of criticality across multiple studies [43, 46, 51]. Compared to young adults, the blood oxygenation level-dependent (BOLD) fMRI timeseries and model dynamics of the neuronal network in the elderly consistently show a lower AC1 (Welch’s t-test, fMRI: *t* = 3.49, *p* = 0.008; Modeling: *t* = 18.90, *p* < 0.0001; see Figure 8**A**). Since the peak in AC1 occurs at criticality, the lower AC1 values observed in older adults suggest that their brain dynamics are more subcritical compared to those of young adults (Figure 8**B**).

**Fig. 8.**
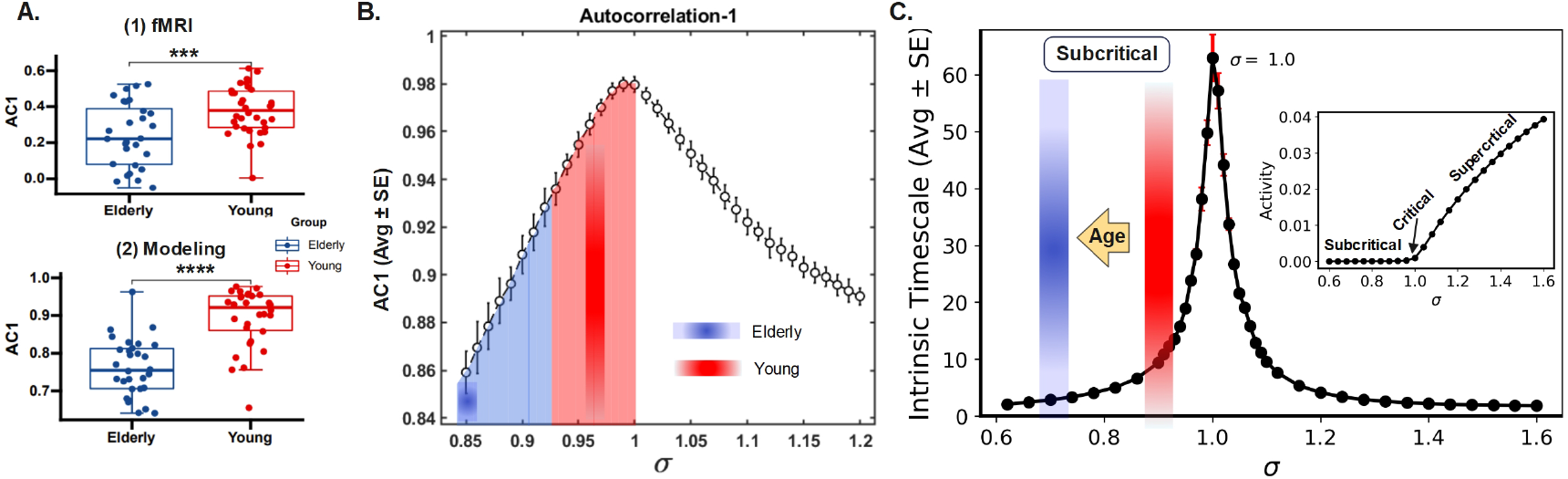
Brain networks in elderly subjects show a greater distance to criticality. **A**. AC1 for fMRI BOLD timeseries (top), and for the modeling dynamics of the neuronal network (bottom). **B**. AC1 peaks at a critical branching ratio (*σ*_*c*_ = 1). The error bar indicates the Standard Error of the Mean over 50 trials. **C**. Intrinsic timescales peak at criticality (*σ*_*c*_ = 1). Advancing age (yellow arrow) shifts the neuronal network towards a further subcritical state. Inset: Neuronal network has a phase transition at a critical branching ratio *σ*_*c*_ = 1; the network is subcritical when the *σ* < 1 and supercritical when *σ* > 1. Activity represents an average across 100 trials.

Finally, we examined the direct relationship between intrinsic timescales and DTC using modeling dynamics. As illustrated in Figure 8**C**, the longest intrinsic timescale of the neuronal network occurs precisely at the critical point (*σ*_*c*_ = 1), where the phase transition occurs. In the subcritical regime, intrinsic timescales increase with the branching ratio. Since the elderly cohort shows a greater DTC compared to the young cohort Figure 8**A**), we can conclude that advancing age shifts the network dynamics further from the critical point, resulting in decreased intrinsic timescales in older adults.

In summary, according to this framework, DTC determines the length of intrinsic timescales, with longer intrinsic timescales occurring near the critical point. Reduced GMV associated with aging leads to a reduced branching ratio in neuronal networks. In turn, this results in networks in the elderly being farther from the critical state, which yields shorter intrinsic timescales. Overall, the aging process, characterized by diminished neuronal connectivity, leads to a more subcritical dynamics (Figure 8**B&C**). This subcritical regime is characterized by enhanced stability but also decreased efficiency in information processing and integration. Consequently, cognitive functions dependent on longer intrinsic timescales, such as memory and executive functions, may experience greater impairment with age.

## 3 Discussion

In this study, we examined changes in intrinsic timescales with age, comparing older adults to young adults, and explained these differences through the dynamics of parsimonious neuronal networks. We observed a hierarchy of intrinsic timescales along the subcortical-cortical axis in both age groups, with shorter timescales in subcortical networks and longer timescales in cortical networks. Importantly, age was associated with decreased intrinsic timescales, and longer intrinsic timescales in the cuneus correlated with better mnemonic discrimination ability (MDA). Moreover, both intrinsic timescales and MDA were positively associated with gray matter volume (GMV), suggesting a neuroanatomical basis for brain function and behavior.

Aiming at further understanding these relationships, age-dependent computational modeling was developed. This approach revealed that alterations in network structure in the aged brain can result in reduced intrinsic timescales. Within the framework of criticality, these age-related changes in intrinsic timescales can be understood through the concept of critical slowing down, where long-range temporal correlations increase as a system approaches a critical point [43–47]. By combining empirical results and neuron network modeling, this study puts forward the idea that altered network properties due to aging increase the distance to criticality, explaining the reduced intrinsic timescales observed with advanced aging. These mechanistic insights enhance our understanding of aging and its effects on brain dynamics and behavior, offering potential pathways for therapeutic interventions.

### 3.1 Criticality Can Explain Alterations in Temporal Dynamics

We explore the mechanism of age-related change of intrinsic timescales through modeling neuronal networks within a potential neuroanatomic basis of GMV. Despite the consistency between our findings and prior work involving the association between GMV and intrinsic timescales, it is essential to note that previous studies did not provide direct evidence to examine how a reduction in GMV may induce changes in intrinsic timescales [7, 27, 29, 31, 35]. Here, our study explores an influential branching process model [53] to gain insight into the neuronal network dynamics representing brain regions for different aging stages, assuming that the GMV reduction observed with age is associated with the pruning of neurons and synapses [14, 16]. We offer evidence that the neuronal networks in young adults are closer to criticality, which is accompanied by longer intrinsic timescales.

Criticality and branching process theory help explain the reduced intrinsic timescales observed with advancing age [49]. Network dynamics can be described effectively by branching processes where there is a critical transition between a low-activity *subcritical* phase and a high-activity *supercritical* phase (Figure 8). This theoretical framework suggests that cortical networks operate near a critical point, where the branching ratio *σ* ≈ 1 [42, 66–68]. At a critical point, the brain optimizes its computational capabilities, responds efficiently to stimuli, and achieves maximal susceptibility, dynamic range, and information processing and storage capacities [41, 42, 49, 50].

Our analysis, using branching process theory, indicates that intrinsic timescales and AC1 are directly dependent on critical transitions [22, 56], peaking at the critical point (*σ*_*c*_ = 1, Figure 8). Although we mainly focused on slightly subcritical neuronal networks, reflecting quasicriticality [56, 64, 69, 70], neuronal networks of young adults are closer to criticality, exhibiting longer intrinsic timescales. In contrast, neuronal networks of elderly subjects are more subcritical, resulting in shorter intrinsic timescales due to age-related reduction in synaptic connectivity [9, 61, 62, 71]. Subcritical neuronal dynamics offer benefits, such as protection from detrimental runaway processes and enhanced specificity in firing-rate coding [48, 63, 72–74]. However, this shift away from criticality also reduces computational capabilities that are maximized at criticality [41, 42, 49, 50].

### 3.2 Brain Criticality with Aging

Using empirical fMRI and neuronal network modeling, our findings provide evidence that the increased distance to criticality accounts for the reduced intrinsic timescales observed in elderly subjects. These findings contribute to the ongoing debate regarding how healthy aging influences brain activity in the context of criticality [49, 75–78]. While a recent study using magnetoencephalography (MEG) data within a quasicriticality framework [78], indicates that aging increases susceptibility, suggesting that the brain is approaching a critical point [79, 80], electroencephalography (EEG) studies across various frequency bands present conflicting evidence [75–77]. These mixed results highlight the complexity of interpreting how criticality evolves with age. Our work, exploring properties of slower low-frequency fluctuations in fMRI BOLD signals, offers a novel perspective on how intrinsic timescales and proximity to criticality evolve with aging. We focused on intrinsic timescales and the autocorrelation function, given their relevance in detecting early-warning signals for critical transitions [43, 44] and critical slowing down [45–47]. Future research should investigate how these metrics relate to other commonly used MEG/EEG measures to further understand the impact of aging on brain dynamics [50]. Additionally, integrating multimodal imaging techniques and advanced computational models should provide deeper insights into the intricate relationship between aging and criticality.

### 3.3 Brain Variability and Model Predictions

The variability of neuronal activity declines with aging, impacting cognitive flexibility [81]. High neural variability supports cognitive flexibility [82] and is a hallmark of critical systems [52]. Furthermore, flexibility decreases as neural systems move from criticality to a subcritical state. A dynamical state with maximum variability of neural patterns occurs at an intermediate coupling strength, and deviations from this optimal point reduce variability [83]. Healthy elderly subjects exhibit a more extensive repertoire of neural activity compared to age-matched subjects with Parkinson’s disease [84] and amyotrophic lateral sclerosis [85]. These findings emphasize the importance of maintaining neuronal variability and flexibility [82], which is optimally achieved near a critical point [52]. Consistent with existing literature [81, 82], our findings of increased subcritical activity in the elderly suggest reduced neural variability. However, this hypothesis should be specifically tested in future studies.

In our age-dependent modeling approach, we simulated normal aging by removing neurons and synapses. Our model indicates that synapse removal increases the distance to criticality and reduces intrinsic timescales, while modest neuron removal, without altering connectivity, has minimal impact. The model highlights the critical role of synapses, as the branching ratio relies only on the connectivity and the mean probability of spiking propagation [53]. This suggests an enhanced importance of synaptic connections over neurons in fundamental network operations [86, 87]. Although our approach involves simplified assumptions, such as treating synaptic weights as static during the brief period used to compute intrinsic timescales, it effectively distinguishes the roles of neurons and synapses. This distinction suggests that removing synapses should result in more subcritical dynamics, leading to reduced intrinsic timescales and decreased number of metastable states or variability in neuronal activity.

### 3.4 Association Between Intrinsic Timescales and Behavior

Intrinsic timescales have been associated with various cognitive functions in both nonhuman human primates, such as delay discounting [19], perception [88, 89], decision making [90] and in humans, such as consciousness [91], the sense of self [92, 93], emotion drawing [94], and visual perception [95]. MDA measures the capability of the brain to distinguish between similar but not identical subjects, which is a crucial high-level cognitive function in daily human activities [58, 96]. Since MDA involves encoding and retrieving information over time [97], its connection to intrinsic timescales seems logical and expected.

We found a positive correlation between MDA and intrinsic timescales at the cuneus during a visual discrimination task, with the cuneus showing enhanced activity during memory retrieval tests [98]. In contrast, no significant correlation was observed between MDA and intrinsic timescales at the whole-brain level, highlighting the specialized role of the cuneus for this memory task. Our results suggest that the shortened autocorrelation of brain activity at the cuneus is closely related to reduced MDA. These findings reinforce the significance of temporal dynamics in cognitive functions, especially within the context of age-related cognitive decline [8, 99, 100].

### 3.5 Intervention

Based on the results of our neuron network modeling, we propose that increasing the excitability of neurons could help *rejuvenate* brain activity in elderly individuals. Our findings indicate that elderly brains have shorter intrinsic timescales due to more subcritical dynamics (*σ* < 1). The model indicates that both network connectivity (*K*) and neuronal excitability (*λ*) influence the distance to criticality (as *σ* = *K* · *λ*). Hence, two types of intervention can be devised based on manipulating either *K* or *λ*. While increasing connectivity (*K*) may be challenging in practical terms, enhancing neuronal excitability (*λ*) is a more feasible approach to bring the branching ratio closer to the critical value (*σ*_*c*_ = 1). By enhancing excitability, brain dynamics could shift closer to the critical state, increasing intrinsic timescales and potentially improving cognitive functions. This intervention aligns with previous proposals that manipulating potassium channels [101] or increasing neuronal excitability [14] could partially restore the physiological integrity of neurons and counteract age-related reductions in neuronal firing rate and dynamic range. However, while longer intrinsic timescales may benefit specific tasks such as memory, it is crucial to consider the potential unintended effects on other brain functions. Shorter intrinsic timescales in the elderly may act as a compensatory mechanism for overall slower brain dynamics that occur with advancing age [102]. Thus, any intervention aimed at modulating intrinsic timescales [40, 103, 104] should be approached cautiously, ensuring a comprehensive understanding of its broader impact on brain activity and cognitive performance.

### 3.6 Hierarchy of Intrinsic Timescales in the Human Brain

We found that the brain functional networks differ in intrinsic timescales, forming a hierarchical topography along the subcortical-cortical networks. The results are consistent with previous studies with fMRI [27, 37–39, 105], which demonstrates that higher-order cortical networks such as the central-executive networks (CEN), dorsal attention networks (DAN), and default-mode network (DMN) have longer intrinsic timescales than sensory and motor regions/networks. This observation of a hierarchy of intrinsic timescales also reflects the function specificity in the brain [19, 26, 97, 103], which is consistent with previous reports about the brain in mice [106], macaque [107– 109], and humans [23, 40, 97, 110]. Our findings reveal that adults, irrespective of age, maintain similarly structured intrinsic timescales, indicating that the hierarchical organization of intrinsic timescales is preserved throughout adulthood. This contrasts with human neonates, who exhibit highly structured intrinsic timescales with a different hierarchy across brain regions that evolve during development [25, 111].

### 3.7 Gray Matter Deterioration as a Neurosubstrate for Reduced Intrinsic Timescales

The decline of intrinsic timescales and MDA with advancing age reflects a reduction in brain functional flexibility [82]. Age-related loss of gray matter volume affects the entire cortex [112–114], suggesting anatomical bases for these declines [99, 115]. The integrity of brain structural networks is crucial for cognitive functions [116], as seen in regions like the hippocampus, which are vital for mnemonic discrimination [96]. Gray matter contains neuronal cell bodies where most neural processing occurs [60]. Reduced gray matter volume means fewer neural resources [60], leading to shorter intrinsic timescales as the brain adjusts to compensate for decreased capacity. Our findings, consistent with studies on autism [27] and Parkinson’s disease [35], show a strong association between gray matter volume and intrinsic timescales. Notably, GMV reduction affects large-scale networks (Figure 6), suggesting that the effects of aging occur in a network-dependent manner rather than in isolated brain areas [117]. This significant correlation between GMV and intrinsic timescales at the network level suggests that declines in GMV contribute to shorter intrinsic timescales by reducing the neuronal network connectivity, thereby making its dynamics more subcritical.

### 3.8 Limitations

Further improvements can be made to our study. Firstly, the age gap between the two cohorts is significant, with young adults averaging 22.21 years and older adults averaging 69.82 years. Previous research has shown that young adults have shorter intrinsic timescales compared to infants, with an average age of 29 years [25, 111]. Another study has reported that older adults (average age 32.8 ± 12.0 years) exhibit shorter intrinsic timescales at specific brain regions, such as the bilateral sensorimotor and occipital regions and the ipsilateral temporal lobe [31]. And some brain regions like the parahippocampal gyrus and putamen show an inverted U-shape relationship with age, increasing in volume until around 45–50 years before decreasing [112], similarly to other brain charts [16]. Taken together, this body of literature suggests that intrinsic timescales may not follow a strictly linear trend across the lifespan, with the dynamics of brain region potentially exhibiting more complex patterns as age progresses.

Additionally, our investigation of GMV utilized peak coordinates derived from independent components analysis rather than specific brain parcellation atlases [118– 120]. These peak coordinates reflect spatial patterns of brain activity rather than direct measures of GMV. As such, they may not capture small-scale alterations in GMV accurately, particularly in complex anatomical regions or overlapping functional networks, potentially leading to underestimation or oversimplification of structural changes. Nevertheless, we observed a robust correlation between GMV and intrinsic timescales in functional regions, highlighting the anatomical basis of brain function dynamics and underscoring the need for further investigation into neuronal network changes.

Lastly, while our current network model provides valuable insights into the relationship between intrinsic timescales and healthy aging, it comprises neurons with stereotypical excitable dynamics. This approach provides a clear view of critical branching processes [53], which was effectively explored here, but may overlook more complex biological phenomena for the sake of clarity. A thorough understanding of brain function will likely require integrating contributions from multiple regions and elements across various organizational levels [121], including neuronal spines [24], dendrites [122, 123], and connectome topology [20, 107].Further research is needed to comprehensively map and incorporate the full range of mechanisms underlying the hierarchy of timescales in the brain and how they evolve throughout the lifespan.

## 4 Methods

Resting-state fMRI scans were obtained from the available public dataset of the University of North Carolina samples at Greensboro [58]. The computational neuronal network was modeled as random networks of excitable spiking neurons. Aging in these neuronal networks was simulated by removing both neurons and synaptic connections. The neuronal network modeling and post-data analysis, including statistical analyses, were carried out with customized code, which is available on [github](https://github.com/xiaohajiayouo/Mapping_and_Modeling_Age-Related_Changes-in-_Intrinsic_Neural_Timescales). All aspects of this study were approved by the Monash University Human Research Ethics Committee.

### 4.1 MRI Data Acquisition, Preprocessing and Head Motion Control

The participants included 28 healthy elderly adults (61–80 years old, mean: 69.82, SD:5.64) and 34 healthy young adults (18–32 years old, mean: 22.21, SD: 3.65). MRI scans include high-resolution T1 anatomical images (1 mm isotropic voxels, 192 slices with thickness of 1mm, repetition time (TR) = 2300 ms, and Time of echo (TE) = 2.26 ms), and functional MRI (32 slices with 4.0 mm thickness and no skip, TE = 30 ms; TR = 2000 ms; flip angle = 70°, field of view = 220 mm, matrix size = 74 × 74 × 32 voxels). The fMRI scan lasted for 10 minutes, which resulted in 300 volumes. Additional details about the raw MRI data can also be found [here](https://openneuro.org/datasets/ds003871/versions/1.0.2).

The first five volumes of each scan were discarded to generate a steady BOLD activity signal to allow for magnetic stability. Similarly to previous studies [124–126], the functional data were then processed with the following steps: (1) Realignment to correct head motion for verification details; (2) slice time correction; (3) outlier identification; (4) normalization (normalize to 3 mm MNI space using a template from the SPM software package [127]); (5) spatial smoothing with a Gaussian kernel of 8 mm full-width at half-maximum (FWHM).

Head motion effects were controlled by calculating the individual framewise displacements (FD). As the outlier scans have been removed from the raw data, none of the remaining participants had head motion greater than 0.5 mm. No significant difference in FD was observed between the two groups FD (*p* = 0.92).

### 4.2 Group ICA for Brain Network Parcellation

A spatial group ICA was performed on the preprocessed and denoised BOLD signal using the Group ICA of FMRI Toolbox (GIFT) infomax algorithm (see Figure 1) [128]. High-model order ICA with a set of 100 components decomposed images into brain regions that comprise larger brain networks (RSN, resting-state functional networks) spanning cortical, subcortical, and cerebellar brain regions. The group ICA back-reconstruction algorithm generated subject-specific spatial maps and time courses from each independent component of the group ICA. After the removal of noise-related components (head motion artifacts, white matter, cerebral spinal fluid, etc.) by visual inspection and regression analysis, we retained 60 components, which represent 60 recognized RSNs. The time courses of non-noise components were triple detrended, despiked, and global average regressed for postprocessing, and finally band-pass filtered (0.023–0.1 Hz) [129].

### 4.3 Gray Matter Volume

Voxel-based morphometry (VBM) preprocessing of T1-weighted structural data was carried out using the Computational Anatomy Toolbox (CAT12) [130]. Specifically, the detailed preprocessing includes the following steps: (1) T1-weighted images were inputted into the segment routine with initial denoising and center-of-mass correction for the gray matter (GM), white matter (WM), and non-brain voxels (cerebrospinal fluid, skull). (2) The population templates (GM, WM) were generated from each dataset separately using the DARTEL algorithm to obtain the corresponding maximum probability map [131]. (3) The gray matter images were aligned to a nonlinear deformation field, and then the normalization parameters were estimated to transform them to MNI space. (4) The normalized images were smoothed according to an isotropic Gaussian kernel (FWHM, full width at half-maximum = 8 mm). The spatially normalized, smoothed, and Jacobian scaled gray matter images (voxel size: 1.5 × 1.5 ×1.5 mm, each image had 131× 137×131, 2,351,057 voxels) were obtained for each participant. (5) the RSN-wise GM and WM were extracted with the coordinate of ICA components. The GM and WM of voxels in the sphere where the centre is the peak coordinate and radius equals 5 mm were averaged. Regions with smaller radii of 3 mm were also tested to assess size robustness. The value represented the final gray matter and white matter volumes of corresponding RSN.

Since voxels with low values of gray matter have variances close to zero and do not apply to the Gaussian error distributions, we masked those low-valued voxels with the [SPM Masking Toolbox](http://www0.cs.ucl.ac.uk/staff/g.ridgway/masking/) with the standard threshold of 0.22 calculated by SPM. As a result, each retained image had a total of 433,584 voxels for further statistical analysis.

### 4.4 Neuronal Network Modeling

#### 4.4.1 Network Topology

Erdös–Rényi undirected random graphs with nodes *N* = 100, 000 and average degree *K* = 10 were utilized to initialize neuronal networks representing a brain region (within the gray matter volume). To simulate healthy aging–characterized by reduced neuronal density [60, 61] and fewer synaptic connections [62, 132, 133] due to neuronal death [134] and decreased neurogenesis [135]–we progressively removed nodes and connections from the network.

Let *R*_*ξ*_ and *R*_*ϕ*_ represent the proportion of *removed* neurons and connections, respectively, of the neuronal network. We assume that both *R*_*ξ*_ and *R*_*ϕ*_ follow normal distributions where 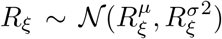 and 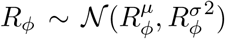, and we set the proportion of removed neurons to be age-dependent following a function based on the empirical measures of GMV:

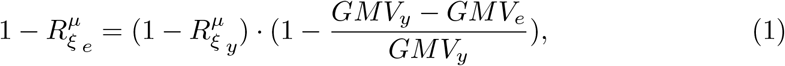

with 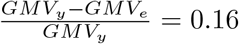, and 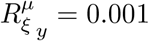, where the subscripts _*y*_ and _*e*_ refer to brain regions in the young and elderly cohorts. Thus, the final mean number of neurons in a network for the two cohorts is 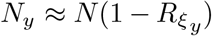 and 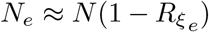. Furthermore, as the variance of GMV between two cohorts is not significant, we set the variance of the 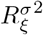 function to be the same for the two groups: 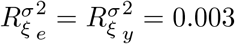.

Likewise, we also consider heterogeneity in the mean number of synaptic connections. To incorporate these synaptic differences, we assumed that the mean number of removed connections for the two cohorts are:

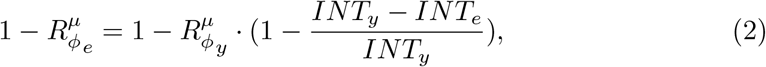

with 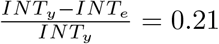 and 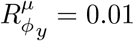. We also assumed the variance in the number of synapses is given by 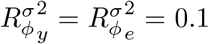. The final mean number of connections *N*_*K*_ for the two cohorts are 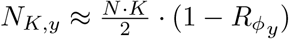, and 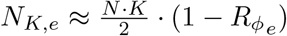.

#### 4.4.2 Neuronal Dynamics

The Kinouchi & Copelli model [53] was used to simulate the dynamics of the neuronal networks. Neuronal networks representing a brain region for young and elderly adults comprised 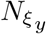 and 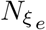 coupled excitable spiking neurons respectively. The adjacency matrix *A* was described by a symmetric binary matrix in which *A*_*ij*_ = *A*_*ji*_ = 1 when neurons *i* and *j* are connected neighbors and *A*_*ij*_ = *A*_*ji*_ = 0 otherwise. The average network degree is given by 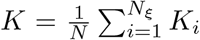, where *K*_*i*_ = *A*_*ij*_ is the node degree. Each neuron *i* is a cyclic cellular automaton that follows a discrete-time process defined on the state space. Specifically, neurons are transiting between 3 states *x*_*i*_: susceptible (*x*_*i*_ = 0), the active (*x*_*i*_ = 1), and refractory state (*x*_*i*_ *>* 1). The states of neurons are updated at each discrete time step of length *δ*_*t*_ = 1 ms. When neuron *i* is at the susceptible state 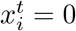, it can be excited either by (i) an active neighbouring neuron 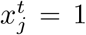 with a fixed probability *λ* = 0.1 if *A*_*ij*_ = 1, or by (ii) an external stimulus which is modelled as a Poisson process with the rate *r* = 10^*−*5^ where *h* = 1 − *exp*^*−r*^. Hence, susceptible neurons are excited with a joint probability *q* = 1 − (1 − *h*) · (1 − *λ*).

An active neuron at time *t* will go to the refractory state at time *t* + 1 and remain there for 8 ms. After that, the neuron will go to the susceptible state. See the flowchart in Figure 1 for a schematic illustration of the neuronal dynamics.

#### 4.4.3 Branching Ratio and Distance to Criticality

The density of active neurons *F* (*t*) of the network was used to compute intrinsic neural timescales of the neuronal network. The network branching ratio is *σ* = *K* · *λ*, and it represents the average number of spikes generated by each excited neuron in the following time step [53]. Critical branching processes occur when *σ*_*c*_ = 1 [41, 50, 64, 65]. For *σ* < 1, the network activity decays over time, indicating a subcritical state. For *σ* > 1, the network activity increases over time, indicating a supercritical state [57]. The branching ratio *σ* is the control parameter of the system, and the distance to criticality for subcritical systems is given by 1 − *σ*.

### 4.5 Autocorrelation Function and Intrinsic Neural Timescales

With the neural activity of brain functional networks and neuronal networks, intrinsic neural timescales were computed based on the autocorrelation function (ACF) of their timeseries, where

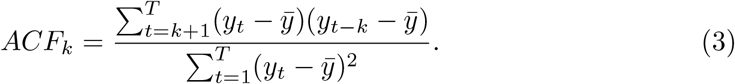

*y* denotes the timeseries of neural activity, 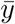 is the mean value across time points, *t* is the time step. For fMRI BOLD timeseries, *t* is the length of time bins, which is the time of repetition (TR = 2000 ms) of the MRI scan, and *T* is the number of time points (295 in the experiment). For the neuronal network timeseries, we set *t* = 1 ms and the *T* = 5050 ms. The first 50 ms was discarded. *k* is the time lag, and a maximum of 25 lag intervals were used.

In most electrophysiological studies, which involve fast dynamics, intrinsic neural timescales (INT) are calculated as the timescale of an exponential decay coefficient fitted to the autocorrelation curve [19, 109, 136]. Here, the intrinsic neural timescales obtained from this fitting method are called (*INT*_*f*_). For resting-state fMRI, which has a lower temporal resolution, the intrinsic neural timescales were computed as the area under the autocorrelation curve to mitigate the adverse effects of low sampling rates [27, 105, 137].

Hence, INT was computed in two different ways for the ICA-recognized functional regions and the neuronal network dynamics. For the functional regions, we calculated the time-dependent magnitude of the autocorrelation function in the resting-state fMRI signals, and intrinsic timescales were computed as the area under the curve (AUC) of the initial ACF curve until it reaches a zero value:

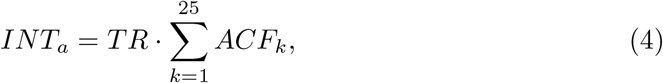

For the neuronal network, we fitted an exponential decay function to the resulting ACF time course using nonlinear least squares, as implemented by the Leven-berg–Marquardt algorithm, using the optimize curve fit function in SciPy. The decay coefficient as the final measure of intrinsic timescales with the following equation [19, 123]:

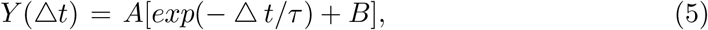

where *τ* represents the intrinsic timescale *INT*_*f*_. *A* is the scaling coefficient, and *B* is the offset, which also represents the asymptotic level of autocorrelation. *INT*_*a*_ were computed for all ICA components, and *INT*_*f*_ were computed for all selected neuronal networks. The exponential fit provides a good approximation for systems sufficiently far from criticality. At criticality, however, the autocorrelation decays more slowly, following a power-law behavior, and the correlation length diverges [42]. In this study, we primarily focus on the subcritical regime, where the exponential fit is fully justified. For consistency and comparison with AC1, we also apply this measure to both critical and supercritical systems.

### 4.6 Reproducibility and Statistical Analysis

The empirical data used in this study are derived from previous research involving the same dataset [58, 124], which includes functional images (fMRI) and T1 weighted images, and the behaviors that involve mnemonic discrimination tasks. These tasks generate a lure discrimination index to evaluate mnemonic discrimination ability (MDA) in both young and elderly participants. To explore the association between intrinsic timescale, MDA, and GMV, the GMV of recognized RSN was measured as the sum of voxels within a sphere where the center is the peak coordinate and the radius equals 5mm. The 3mm was selected as an alternative to examine the robustness of the results. Computational neuronal network modeling was based on parsimonious excitable networks [53]. We simulated neuronal networks across different ages by removing nodes and connections and analyzed 60 randomly selected samples of neuronal networks (30 per cohort) to assess differences in intrinsic timescales with aging. Additional results from larger sample sizes—100 (50 per cohort) and 200 (100 per cohort)—are provided in the supplementary material.

Statistical analyses were conducted using common parametric tests, including Wilcoxon rank-sum tests, t-tests, and F-tests. For significant ANOVA results, post hoc analyses were performed using Welch’s t-test. Multiple comparisons were performed using false discovery rate (FDR) correction with a threshold of *p* < 0.05. All statistical analyses and computational neuronal network modeling were carried out with customized code, which is made available online, as detailed in the “Code availability” section.

## Data Availability

All behavioral and raw MRI data used in this study were from the public [dataset](https://openneuro.org/datasets/ds003871/versions/1.0.2). The timeseries of independent components used to recognize the brain functional networks and the smoothed whole brain GMV can be found in [GitHub Repository](https://github.com/xiaohajiayouo/Mapping_and_Modeling_Age-Related_Changes-in-_Intrinsic_Neural_Timescales) Data folder. In addition, this folder has 200 samples of produced neuronal networks. Each one contains the network dynamics (timeseries), the branching ratio, and the corresponding computed intrinsic timescale. All data present in this manuscript has been sorted out and tidied up into Excel files.

## Code Availability

The code for neuronal networks modelling, post-data analysis including the intrinsic timescale estimation, GMV comparison, and statistics analysis, as well as the result visualization, are all available here [Github](https://github.com/xiaohajiayouo/Mapping_and_Modeling_Age-Related_Changes-in-_Intrinsic_Neural_Timescales).

## Funding

This work was supported by the Australian Research Council (ARC) Future Fellowship (FT200100942) and the Rebecca L. Cooper Foundation (PG2019402).

## Supplemental Data

The supplemental materials and data can be found in the attached Supplementary Materials.

## Notes

### Competing Interest Statement

The authors have declared no competing interest.

