## supplemental Material for "Mapping and Modeling Age-Related Changes in Intrinsic Neural Timescales"

### *Supplementary Material*

Kaichao Wu, Leonardo L. Gollo

**Table I. Peak coordinates of ICNs**

| ICNs | X | Y | Z |
|---|---|---|---|
| **Default-mode Domain: 11** | | | |
| Anterior cingulum cortex (18) | -0.5 | 25.5 | -13.5 |
| Precuneus (19) | -0.5 | 58.5 | 64.5 |
| Anterior cingulum cortex (31) | -0.5 | -34.5 | 4.5 |
| Anterior cingulum cortex (52) | -0.5 | -4.5 | -10.5 |
| Angular gyrus (53) | 47.5 | -64.5 | 37.5 |
| Post cingulum + Precuneus (55) | -0.5 | -43.5 | 43.5 |
| Post cingulum (64) | -0.5 | -28.5 | 26.5 |
| Post cingulum (74) | 14.5 | -46.5 | -1.5 |
| Post cingulum + Precuneus (86) | -0.5 | -62.5 | 19.5 |
| Post cingulum + Precuneus (87) | -0.5 | -64.5 | 37.5 |
| Post cingulum (94) | -0.5 | -46.5 | 4.5 |
| **Sensorimotor Domain: 6** | | | |
| Right precentral gyrus (1) | 35.5 | -19.5 | 67.5 |
| Postcentral gyrus (15) | -57.5 | -7.5 | 25.5 |
| Superior parietal lobule (20) | 23.5 | -43.5 | 67.5 |
| Left precentral gyrus (21) | -36.5 | -22.5 | 61.5 |
| Paracentral lobule (25) | -0.5 | -22.5 | 58.5 |
| Postcentral gyrus (48) | 59.5 | -25.5 | 34.5 |
| **Visual Domain: 12** | | | |
| VIS.Middle occipital gyrus (23) | 29.5 | -91.5 | 1.5 |
| VIS.Lingual gyrus (17) | -0.5 | -82.5 | -7.5 |
| VIS.Cuneus (30) | 2.5 | -85.5 | 10.5 |
| VIS.Cuneus (34) | 2.5 | -76.5 | 43.5 |
| VIS.Middle temporal gyrus (34) | 47.5 | -64.5 | -19.5 |
| VIS.Lingual gyrus (37) | -0.5 | -64.5 | -4.5 |
| VIS.Lingual gyrus (47) | -23.5 | -64.5 | -13.5 |
| VIS.Left Lingual gyrus (57) | -24.5 | -76.5 | -16.5 |
| VIS.Middle temporal gyrus (58) | 53.5 | -67.5 | 1.5 |
| VIS.Calcarine (67) | 14.5 | -61.5 | 7.5 |
| VIS.Cuneus (79) | 32.5 | -76.5 | 22.5 |
| VIS.Cuneus + precuneus (89) | 26.5 | -64.5 | 49.5 |
| **Cognitive-control Domain: 18** | | | |
| CC.Supplementary motor area (23) | -0.5 | -28.5 | -58.5 |
| CC.Left inferior frontal gyrus (28) | -51.5 | 34.5 | -1.5 |
| CC.Middle cingulum (33) | -0.5 | 10.5 | -34.5 |
| CC.Supplementary motor area (35) | -0.5 | -4.5 | 64.5 |
| CC.Left Insula (39) | -36.5 | 7.5 | -25.5 |
| CC.Superior frontal gyrus (42) | -24.5 | 19.5 | 49.5 |
| CC.Insula (43) | -42.5 | 13.5 | -7.5 |
| CC.Left Inferior parietal lobule (46) | -39.5 | -43.5 | 43.5 |
| CC.Left Lingual gyrus (60) | -12.5 | -52.5 | -1.5 |

| | | | |
|---|---|---|---|
| CC.Post cingulum (63) | -3.5 | -37.5 | -7.5 |
| CC.Middle frontal gyrus (65) | -51.5 | 13.5 | 28.5 |
| CC.Superior medial frontal gyrus (68) | -0.5 | 46.5 | 43.5 |
| CC.Insula (70) | -42,5 | -7.5 | -13.5 |
| CC.Right inferior frontal gyrus (73) | 56.5 | 16.5 | 19.5 |
| CC.Middle frontal gyrus (78) | -33.5 | 55.5 | -1.5 |
| CC.Middle frontal gyrus (83) | -0.5 | 52.5 | 19.5 |
| CC.Supplementary motor area (85) | -0.5 | -16.5 | 49.5 |
| CC.Right Inferior parietal lobule (88) | 44.5 | -46.5 | 52.5 |
| **Sub-cortical Domain: 7** | | | |
| SC.Caudate (25) | -0.5 | -4.5 | 16.5 |
| SC.Caudate (40) | -0.5 | 7.5 | 1.5 |
| SC.Putamen (49) | -27.5 | -46.5 | -1.5 |
| SC.Caudate (99) | 5.5 | -1.5 | 10.5 |
| SC.Caudate (66) | 8.5 | -19.5 | 19.5 |
| SC.Posterior thalamus (76) | -0.5 | -22.5 | 7.5 |
| SC.Medial thalamus (90) | -0.5 | -22.5 | -4.5 |
| **Auditory Domain: 6** | | | |
| AUD.Superior temporal gyrus (69) | -54.5 | -25.5 | 10.5 |
| AUD.Superior temporal gyrus (75) | -57.5 | -37.5 | -4.5 |
| AUD.Superior temporal gyrus (77) | -60.5 | -1.5 | -1.5 |
| AUD.Right Superior temporal gyrus (91) | 44.5 | -7.5 | -1.5 |
| AUD.Right Superior temporal gyrus (96) | 59.5 | -43.5 | 4.5 |
| AUD.Left Superior temporal gyrus (97) | -54.5 | -58.5 | 16.5 |

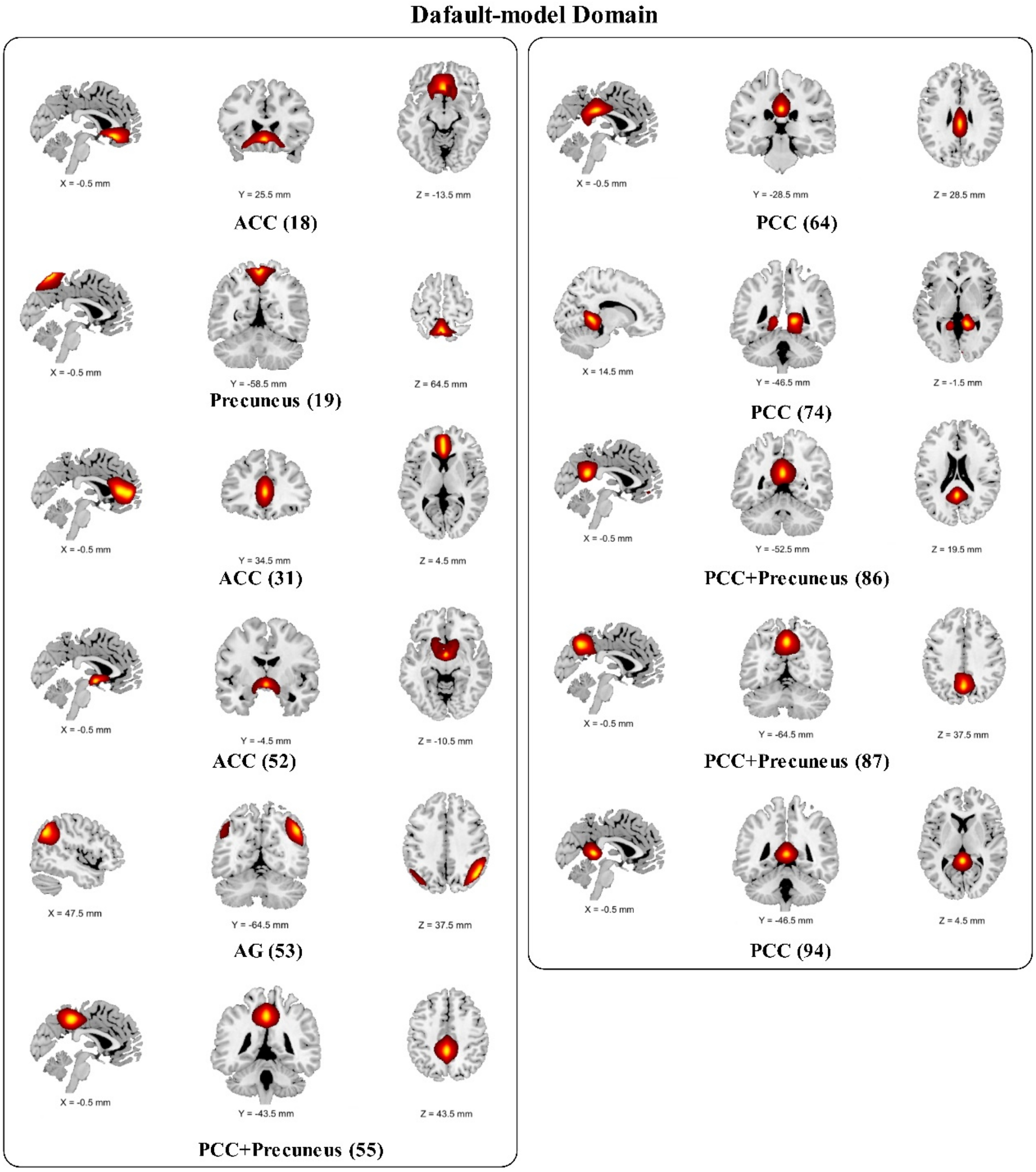


**Figure 1-1. Spatial maps of intrinsic connectivity networks in the default mode domain.** Sagittal, coronal, and axial slices are shown at the maximal t-statistic for clusters larger than 3 $cm^3$.

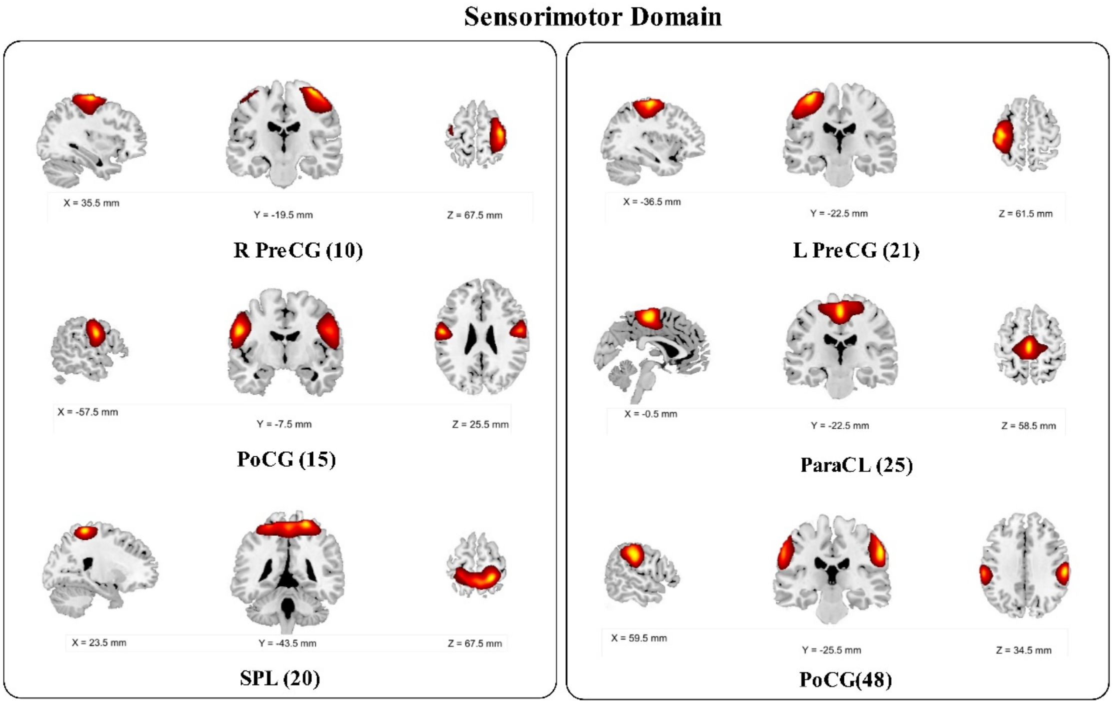


**Figure 1-2. Spatial maps of intrinsic connectivity networks in the sensorimotor domain.** Sagittal, coronal, and axial slices are shown at the maximal t-statistic for clusters larger than 3 $cm^3$.

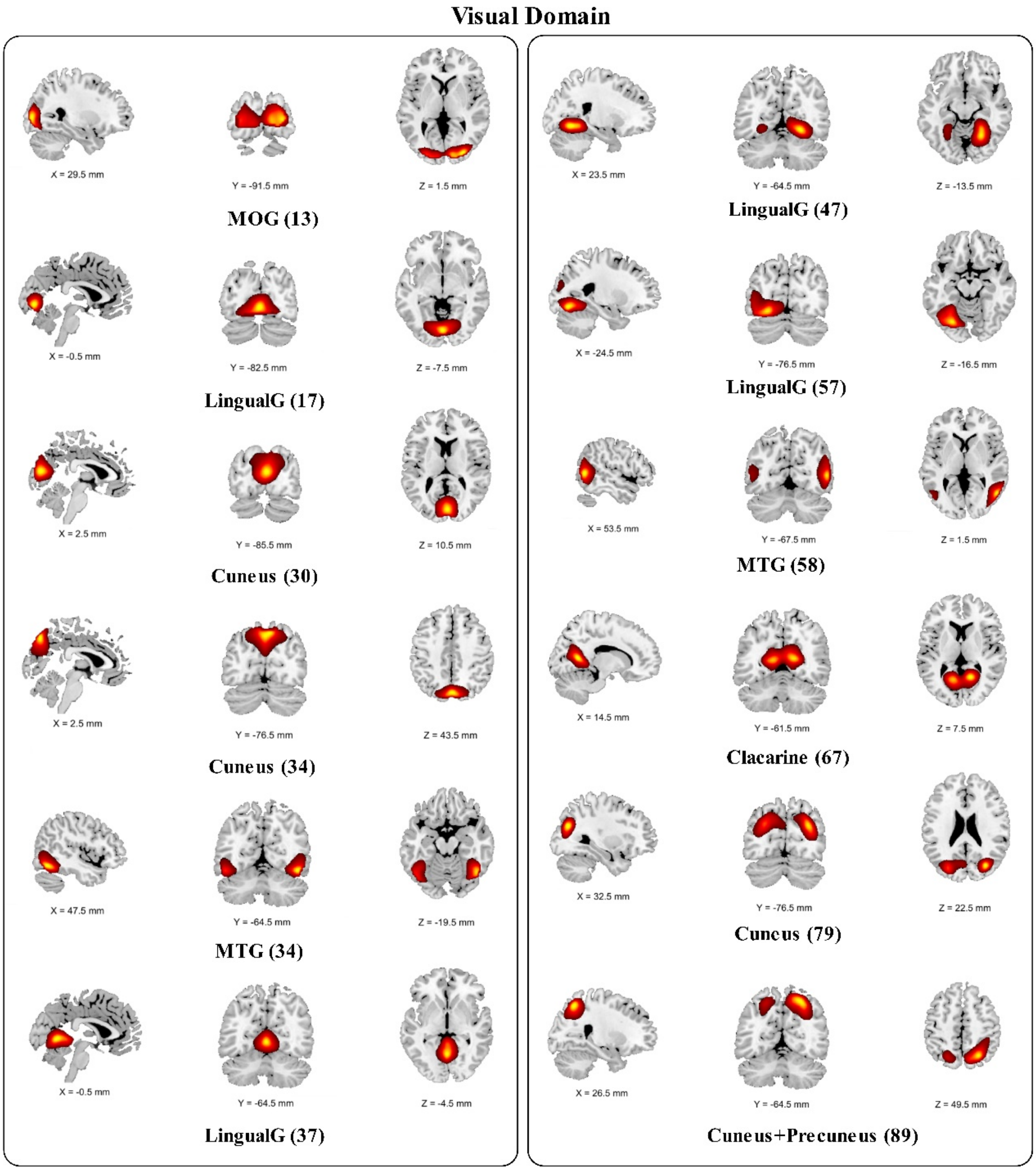


**Figure 1-3. Spatial maps of intrinsic connectivity networks in the visual domain.** Sagittal, coronal, and axial slices are shown at the maximal t-statistic for clusters larger than 3 $cm^3$.

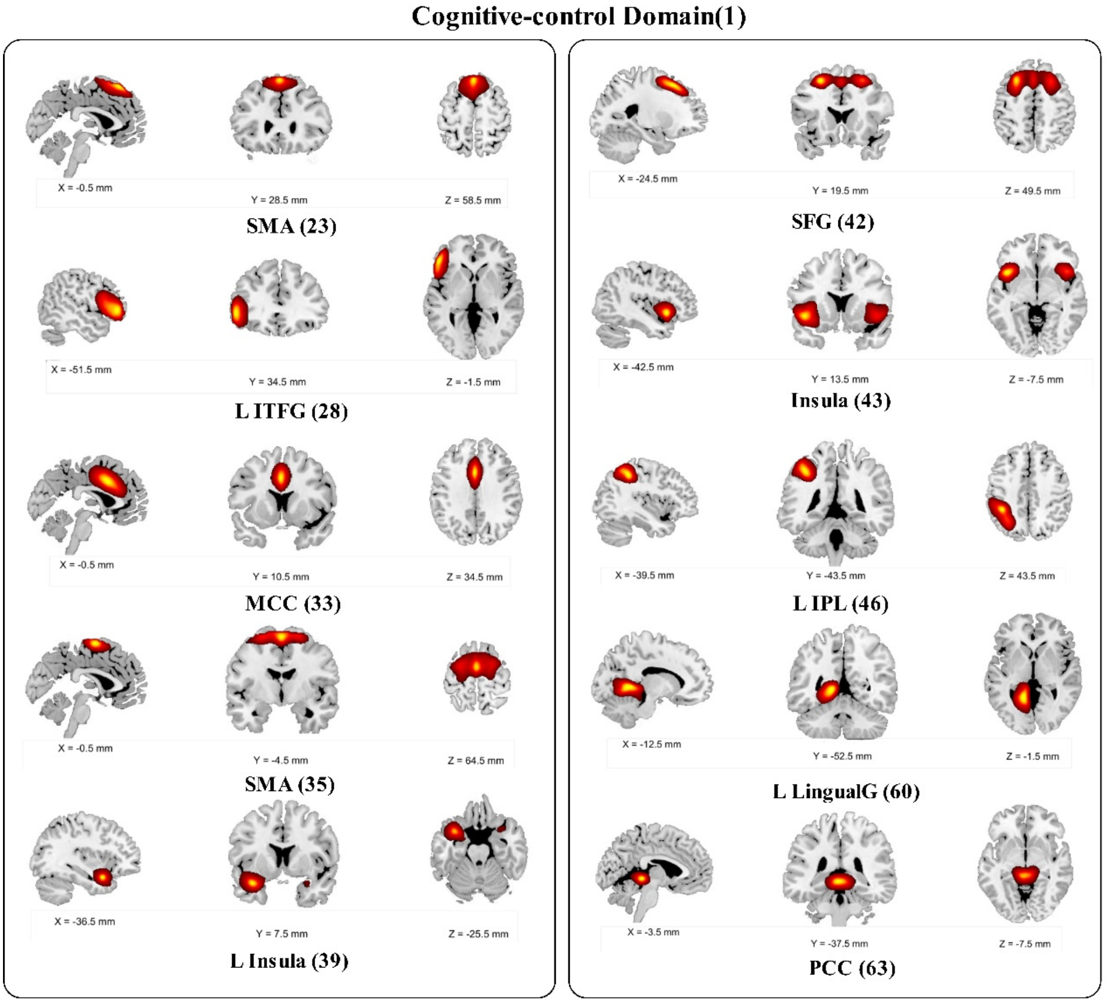


**Figure 1-4(1). Spatial maps of intrinsic connectivity networks in the cognitive control domain.** Sagittal, coronal, and axial slices are shown at the maximal t-statistic for clusters larger than 3 $cm^3$.

**Cognitive-control Domain(2)**

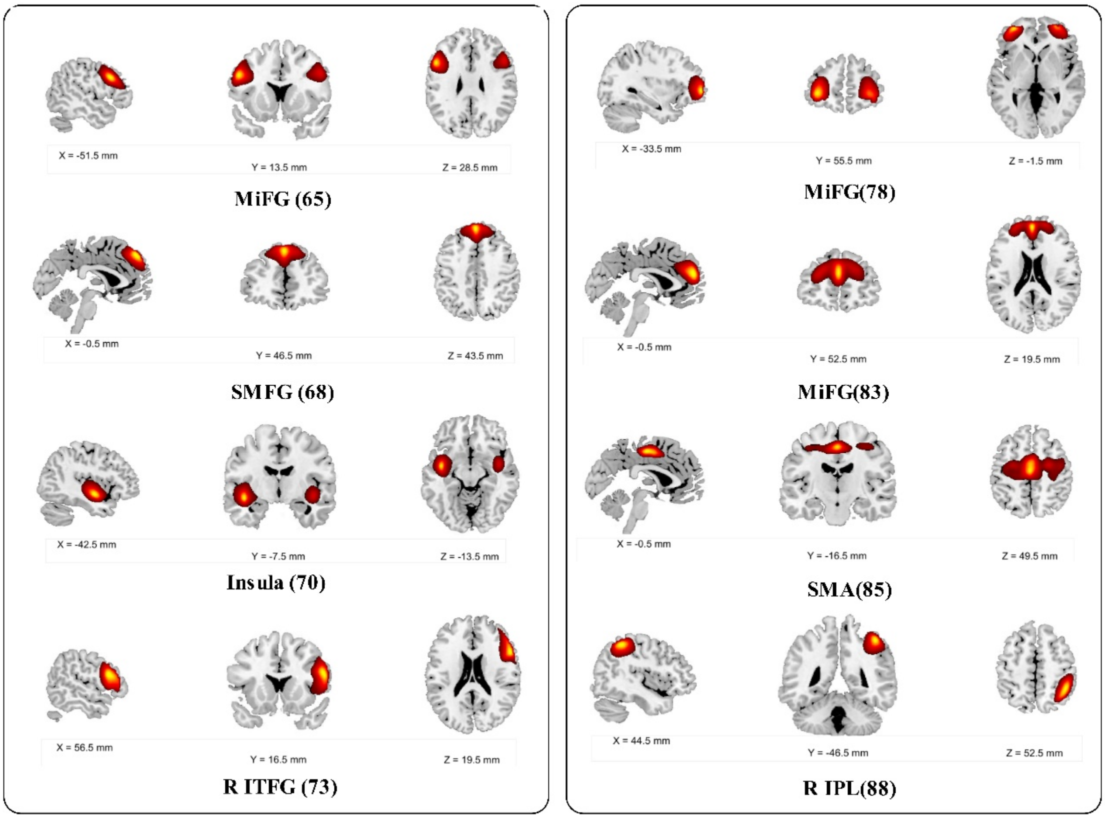


**Figure 1-4(2). Spatial maps of intrinsic connectivity networks in the cognitive control domain.** Sagittal, coronal, and axial slices are shown at the maximal t-statistic for clusters larger than 3 $cm^3$.

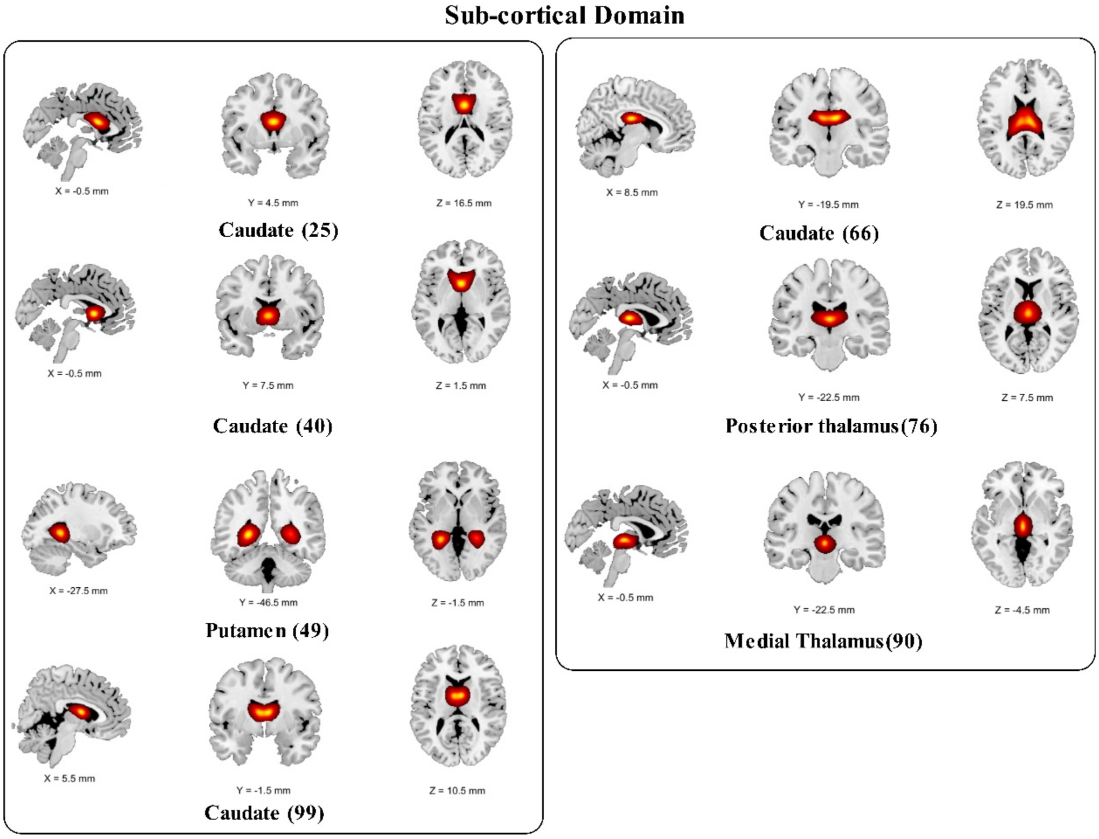


**Figure 1-5. Spatial maps of intrinsic connectivity networks in the subcortical domain.** Sagittal, coronal, and axial slices are shown at the maximal t-statistic for clusters larger than 3 $cm^3$.

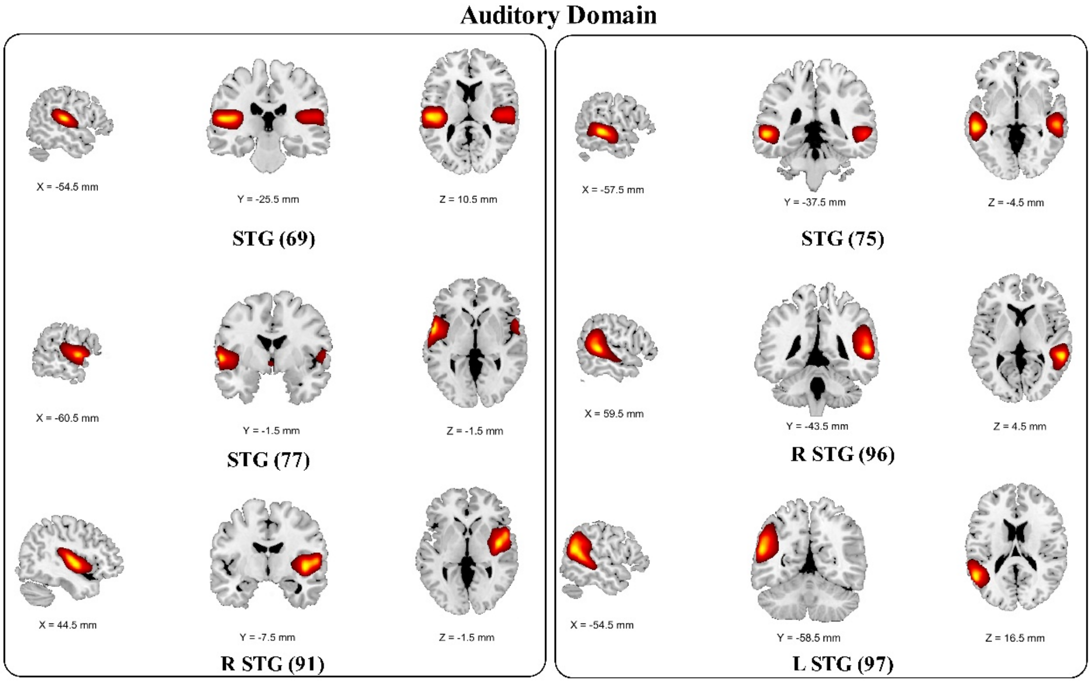


**Figure 1-6. Spatial maps of intrinsic connectivity networks in the subcortical domain.** Sagittal, coronal, and axial slices are shown at the maximal t-statistic for clusters larger than 3 $cm^3$.

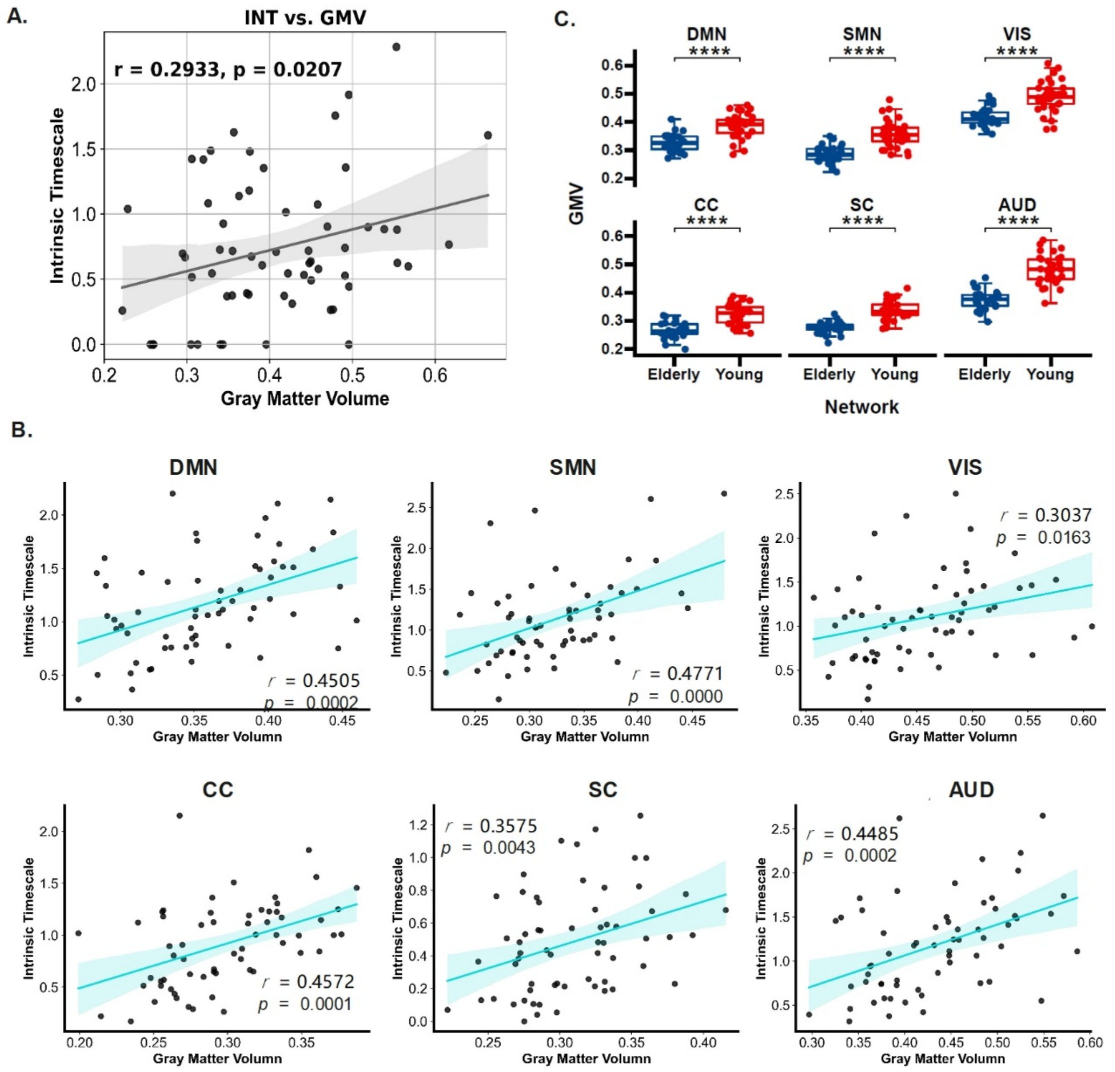


**Figure 2. Robustness in the relationship between GMV and intrinsic timescales.** The relevant result of GMV extracted with a radius = 3mm. **A.** Positive association between GMV and INT of cuneus can be observed. **B**. Network-level comparison of GVM between the elderly and young. **C.** Association between GMV and INT at the network level.

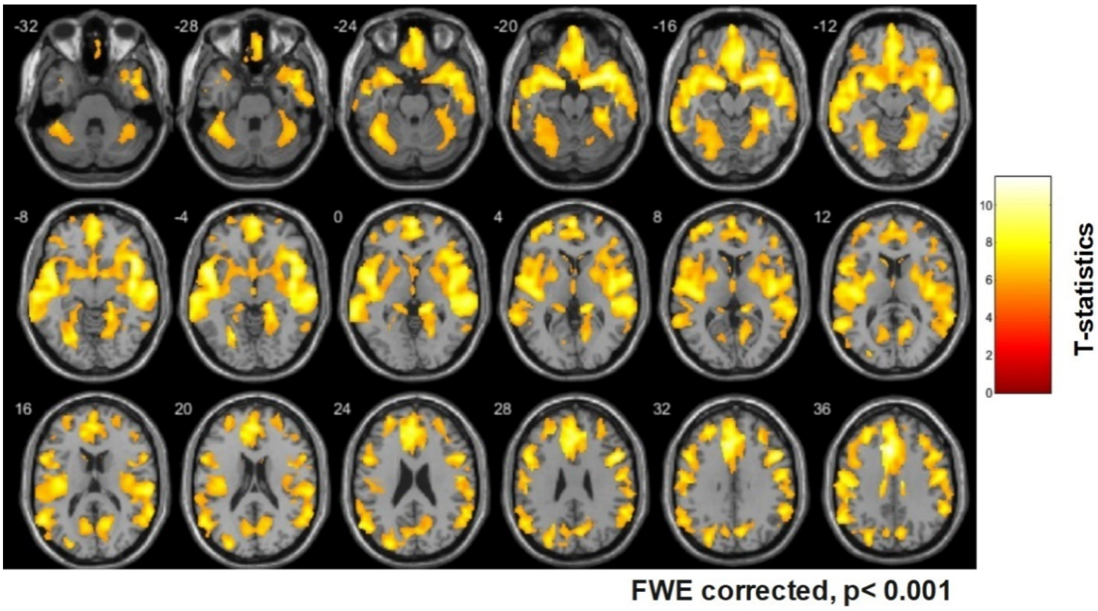


**Figure 3.** The whole brain comparison of GMV between the young and elderly.

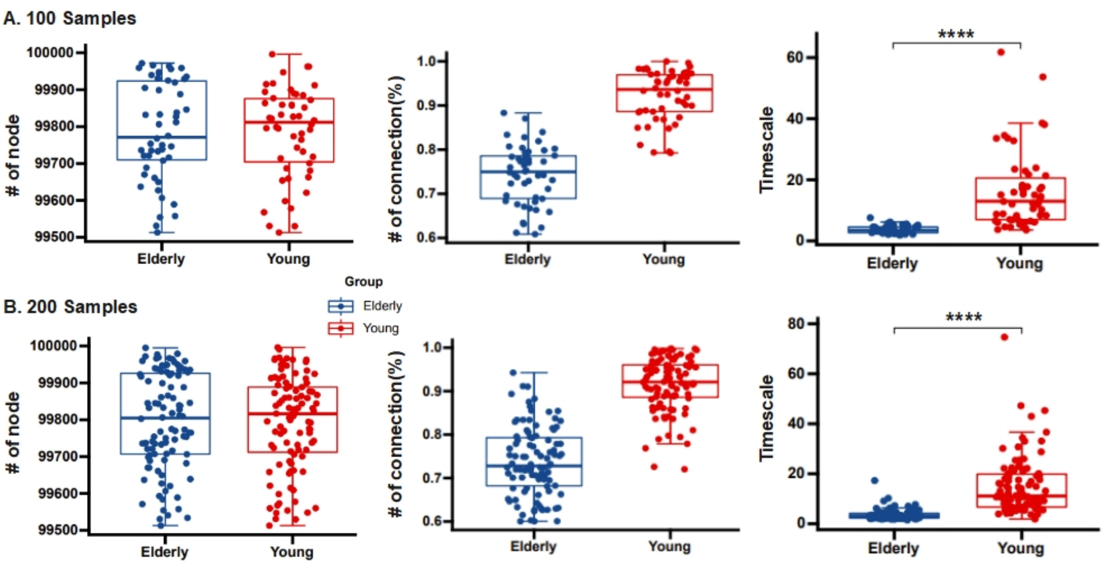


**Figure 4.** Robustness of modeling results. Larger intrinsic timescales can be robustly observed either with (A) 100 samples (50 young and 50 elderly) or (**B**) 200 samples (100 young and 100 elderly).
